# Mediator is broadly recruited to gene promoters via a Tail-independent mechanism

**DOI:** 10.1101/2021.12.21.473728

**Authors:** Linda Warfield, Rafal Donczew, Lakshmi Mahendrawada, Steven Hahn

## Abstract

Mediator (MED) is a conserved factor with important roles in both basal and activated transcription. It is believed that MED plays a direct role in transcriptional regulation at most genes by functionally bridging enhancers and promoters. Here, we investigate the genome-wide roles of yeast MED by rapid depletion of its activator-binding domain (Tail) and monitoring changes in nascent transcription. We find that MED Tail and activator-mediated MED recruitment regulate only a small subset of genes. At most genes, MED bypasses the UAS and is directly recruited to promoters to facilitate transcription initiation. Our results define three classes of genes that differ in PIC assembly pathways and the requirements for MED Tail, SAGA, TFIID and BET factors Bdf1/2. We also find that the depletion of the MED middle module subunit Med7 mimics inactivation of Tail, suggesting a functional link. Our combined results have broad implications for the roles of MED, other coactivators, and mechanisms of transcriptional regulation at different gene classes.

## Introduction

Mediator (MED) is a conserved eukaryotic transcription factor involved in many facets of gene expression with principal functions in both basal and regulated transcription (Allen and Taatjes, 2015; Jeronimo and Robert, 2017; Soutourina, 2018). In most systems, genome-wide RNA Pol II transcription is strongly decreased upon MED inactivation (El_Khattabi et al., 2019; Holstege et al., 1998; Jaeger et al., 2020; Petrenko et al., 2017; Warfield et al., 2017). Mutations in MED subunits can affect many signaling and developmental pathways and, in many cases, have been linked to disease (Allen and Taatjes, 2015; Jeronimo and Robert, 2017; Soutourina, 2018). Structural and biochemical studies found that MED is organized into four structural modules: Head, Middle and Tail, linked together by the connector subunit Med14, and containing a less tightly associated kinase module (Harper and Taatjes, 2018). Head and Middle modules comprise the MED core and contain nearly all the subunits essential for viability while Tail subunits generally have gene-specific roles.

MED has several well-established biochemical functions important for basal transcription including: (i) MED binds Pol II with nM affinity primarily via the CTD (Robinson et al., 2016). In agreement with this, a substantial fraction of MED in yeast cell extracts is bound to Pol II (Kim et al., 1994; Li et al., 1995; Liu et al., 2000). (ii) MED stimulates preinitiation complex (PIC) formation both in vitro and in vivo and MED depletion can lead to severe defects in PIC formation and gene expression (Baek et al., 2006; Eychenne et al., 2016; Nguyen et al., 2021; Petrenko et al., 2017; Ranish et al., 1999). (iii) In vitro, MED stably binds the PIC, interacting with the Pol II CTD, several other Pol II surfaces and with the basal transcription factors TFIIE and TFIIH (Abdella et al., 2021; Chen et al., 2021; Rengachari et al., 2021; Robinson et al., 2016; Schilbach et al., 2017). Although the MED-PIC complex is stable in vitro, yeast MED has not been detected in vivo at promoters during normal growth but instead at a subset of enhancers/UAS elements (Grünberg et al., 2016; Jeronimo and Robert, 2014; Knoll et al., 2018). MED has only been observed at yeast promoters after slowing escape of Pol II from the PIC by depletion or inhibition of the CTD kinase Kin28 (CDK7) (Jeronimo and Robert, 2014; Knoll et al., 2018; Wong et al., 2014). Together, these findings suggested that the MED-PIC complex is short lived and that Pol II initiates transcription soon after formation of MED-PIC. (iv) MED stimulates CTD phosphorylation by CDK7/Kin28 (Kim et al., 1994). CTD phosphorylation is needed for release of Pol II from the MED-PIC complex and structural studies revealed that MED positions CDK7/Cyclin H in the PIC where it faces the CTD and is in position to processively phosphorylate CTD repeats (Abdella et al., 2021; Chen et al., 2021; Rengachari et al., 2021). Finally, the interaction of CDK7 with MED subunit Med6 may activate kinase activity.

While these roles of MED in basal transcription have been well characterized, many questions remain about the role of MED in transcriptional regulation. A consensus view is that MED transduces signals from transcription activators to the transcription machinery by acting as a functional bridge between enhancers and promoters (Allen and Taatjes, 2015; Jeronimo and Robert, 2017; Soutourina, 2018). In support of this model, many transcription activators are known to bind MED subunits that are required for activated transcription. In yeast, most activator-MED interactions occur within the Tail module, comprised of subunits Med2, 3, 5, 15, and 16 (Fishburn et al., 2005; Lee et al., 1999; Park et al., 2000; Reeves and Hahn, 2005; Thakur et al., 2008; Zhang et al., 2004). Yeast Med15 is the best characterized activator target and contains four activator-binding domains (Brzovic et al., 2011; Fishburn et al., 2005; Herbig et al., 2010; Jedidi et al., 2010; Park et al., 2000; Thakur et al., 2008; Tuttle et al., 2021). In mammalian MED, other frequent activator targets include Med1 and Med25; summarized in (Abdella et al., 2021). However, despite decades of work, it’s not clear how activator-MED binding stimulates transcription. An early proposal was that transcription activators, binding at enhancers, recruit MED to gene regulatory regions and thereby facilitate formation of the functional MED-PIC complex via transfer of MED from enhancer to promoter (Ptashne and Gann, 1997; Struhl, 1996). This recruitment model is supported by ‘artificial recruitment’ experiments where linking a MED Tail subunit to a high affinity DNA binding domain can lead to high levels of transcription in the absence of a conventional activator (Barberis et al., 1995; Cheng et al., 2004; Farrell et al., 1996; Jiang and Stillman, 1992; Keaveney and Struhl, 1998). Consistent with the recruitment model, it was reported that a single molecule of MED can crosslink to both promoter and UAS element (Petrenko et al., 2016). A second non mutually exclusive model is that activator-MED binding facilitates a conformational change necessary for Pol II binding (Bernecky and Taatjes, 2012; El_Khattabi et al., 2019; Meyer et al., 2010; Zhang et al., 2021). The relative positioning of the Head, Middle and Med14 are known to change upon binding of MED to Pol II and the PIC (Schilbach et al., 2017; Tsai et al., 2017). Facilitation of this conformational change by activators could conceivably overcome a rate limiting step in transcription. However, all known activator binding domains in MED appear tethered via intrinsically disordered regions so it’s not understood how AD-MED interactions could lead to a specific conformational change.

Despite compelling arguments for MED involvement in activated transcription of some genes, the number of genes regulated by this pathway is unclear. For example, individual yeast Tail subunits are not essential for cell viability and genome-wide studies have observed changes in the steady state mRNA levels for only ∼10% of genes upon inactivation of individual Tail subunits (El_Khattabi et al., 2019; Knoll et al., 2018; Larsson et al., 2013; Myers et al., 1999; Petrenko et al., 2017; van_de_Peppel et al., 2005). Interestingly, both positive and negative changes in expression of individual genes were observed upon Tail inactivation in both yeast and mammalian cells. In contrast, Tail inactivation has been observed to cause broad genome-wide decreases in Pol II ChIP signals and moderate decreases in PIC formation at most genes, suggesting a more general role (Jeronimo et al., 2016; Knoll et al., 2018). Further confounding this issue is that MED has been observed at many core promoters upon CDK7 inhibition, even in strains that lack functional Tail (Jeronimo et al., 2016; Knoll et al., 2018; Petrenko et al., 2016). Together, these findings led to the proposal that MED recruitment via enhancers and the Tail is the dominant pathway but that there is an alternative pathway whereby MED can be recruited directly to core promoters (Jeronimo and Robert, 2017; Jeronimo et al., 2016; Knoll et al., 2018). Determining the significance of this alternative pathway under both normal growth and stress conditions will have important implications for understanding the roles of MED in transcriptional regulation.

Here, we have examined the relative importance of these two MED recruitment pathways for genome-wide transcription. Our results suggest that the MED Tail and direct activator-mediated MED recruitment regulates only a small subset of genes. At most genes, MED bypasses the UAS and is directly recruited to promoters where it has important functions in PIC formation and transcription initiation. Our results define three types of yeast genes having unique PIC assembly pathways and coactivator requirements and they have important implications for the functions of transcription factors, coactivators, and MED at these different gene classes.

## Results

### A small set of Mediator Tail-dependent genes

To identify Tail-dependent genes, we first measured changes in newly synthesized mRNA after rapid degradation of Tail subunits. Subunits of Tail were fused to the auxin degron IAA7 (Chan et al., 2018; Nishimura et al., 2009) and strains grown in synthetic complete media were treated with either DMSO (control) or 3-Indoleacetic acid (IAA) for 30 min. Cells were then briefly labeled with 4-Thiouracil and labeled RNA was isolated and analyzed by RNA-seq (Donczew et al., 2020a). In our initial experiments, we depleted the Tail subunits predicted from prior work to be most important for Tail function: (i) the activator binding subunit Med15, (ii) simultaneous depletion of Med15 and Med2 (Med2 interacts with the Med14 C-terminal domain), and (iii) depletion of the Med14 C-terminal domain that is known to be required for association of Tail with MED core (Lee et al., 1999; Li et al., 1995). For this latter cell line, the N and C-terminal domains of Med14 (Tsai et al., 2017) were expressed on separate plasmids: (a) the Med14 C-terminal region (termed the Tail interaction domain (TID); residues 705-1082) fused to the IAA7 degron and (b) the Med14 N-terminal region (Med14N; residues 1-746). Expression of both these Med14 derivatives was driven by the *MED14* promoter. In the absence of IAA, growth of nearly all degron strains used in this work was similar to WT, with the exception that strains expressing separate Med14N and TID polypeptides grew slightly slower (**Fig S1A**). Protein degradation was assessed by Western analysis where we found ≤10% of the degron-fusion proteins remained after 30 min of IAA treatment (**Fig S1B**). All RNA-seq experiments, unless otherwise noted, were performed in triplicate from cells grown in synthetic complete glucose media and variation between replicate samples was <20% (**Fig S1C**).

Transcription from only a modest number of genes was altered upon rapid Tail subunit depletion with the Med15 and Med14 TID-degron strains showing the strongest transcriptional changes (**Table S1**). K-means clustering was used to sort genes into two categories based on transcriptional changes in the three degron strains (**Fig 1A, B**). Of the 4885 genes that can be reliably quantitated under these growth conditions, we found that only 287 (5.7% of expressed genes) fell into the Tail-dependent category. 90% of these genes showed decreased expression, ranging from 1.3-4.0-fold lower expression, after Med15 or TID inactivation. Further, expression from ∼60% of this gene set (∼170 genes) was reduced ≥ 1.5-fold upon Med15 or TID inactivation (**Fig S2A**). Our results contrast with earlier studies that used gene deletions and steady state mRNA analysis and observed approximately equal numbers of upregulated and downregulated genes upon long-term Tail inactivation (Ansari et al., 2011; El_Khattabi et al., 2019; Knoll et al., 2018). To compare transcription defects caused by rapid depletion vs gene deletions, we measured changes in levels of newly synthesized mRNA in strains containing *med15Δ* or *med16Δ* as well as in strains containing degrons fused to other Tail subunits: Med5, Med16, and Med15+16 (**Fig 1B; Fig S2B**). Comparison of all results showed that the Med15 deletion gave the strongest median transcription loss from Tail-dependent genes, followed closely by the Med15 and Med14 TID-degrons. The Med16-degron strain showed only very modest changes (median change of 1.2-fold for Tail-dependent genes), and the Med5-degron showed little difference from wild type. While Med15 and 16 deletions largely affect transcription from Tail-dependent genes, changes due to the Med15 deletion poorly correlate with results from the Med15 degron (**Fig 1C**; r= 0.2). This difference likely reflects additional indirect effects on transcription due to the long-term absence of Med15 in the deletion strain.

**Fig 1.**
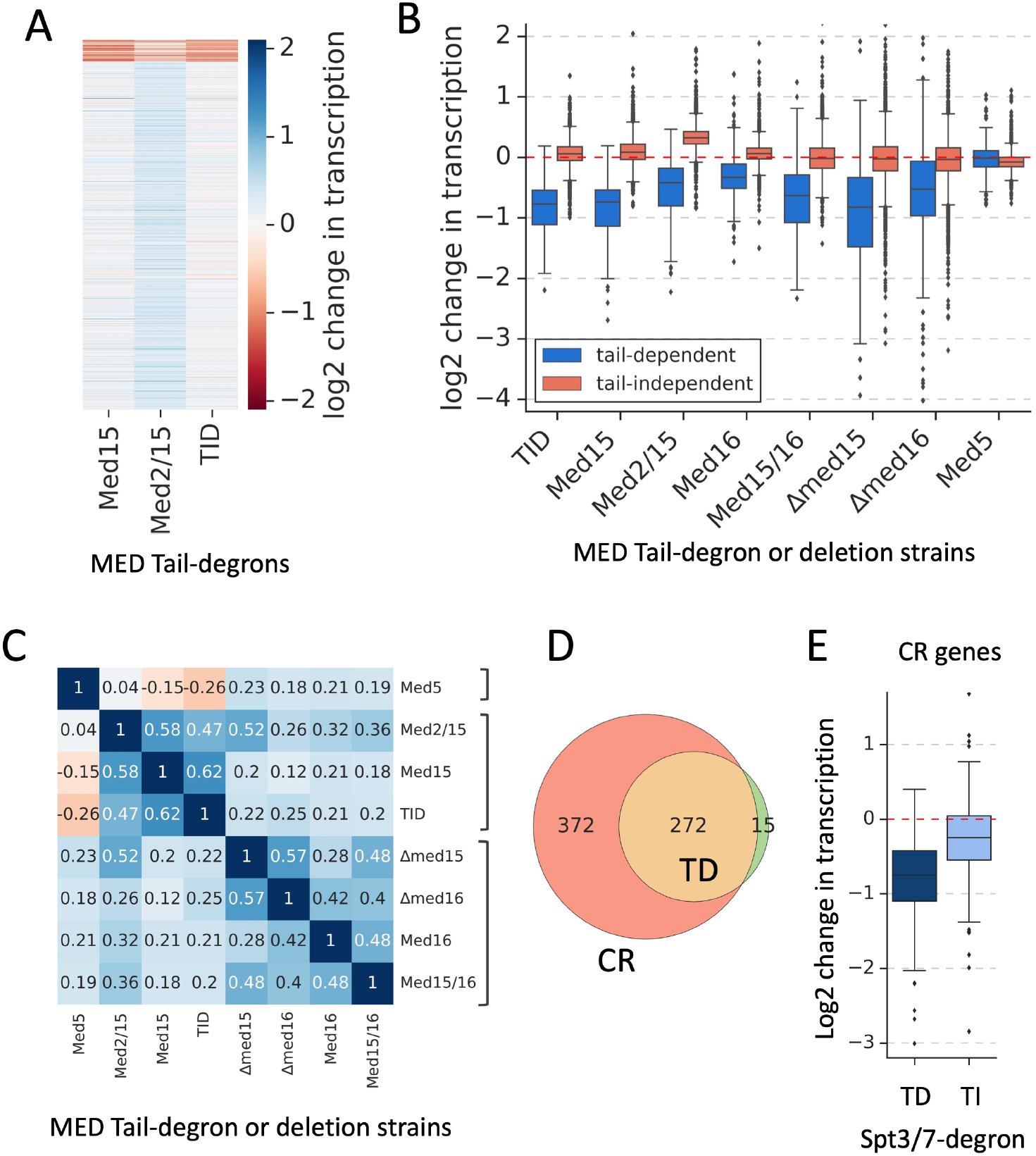
Mediator Tail-dependent genes are a subset of coactivator-redundant (CR) genes. **(A)** Heatmap representation of log2 change in transcription levels upon MED Tail inactivation. Genes are grouped by results of k-means clustering analysis of Mediator Tail-depletion experiments assayed by 4-ThioU RNA-seq. Two clusters were found to give the best separation using silhouette analysis. Log2 change values from relevant experiments for all 4885 genes were used as an input for k-means algorithm (‘KMeans’ function from Python sklearn.cluster library with default settings). **(B)** Boxplot showing log2 change in transcription for 4885 genes after depleting the indicated factors. Genes are grouped into Tail-dependent and Tail-independent categories. **(C)** Matrix of Spearman correlation coefficient values for log2 changes in transcription in the indicated experiments. Brackets indicate correlated groups. **(D)** Venn diagram showing the overlap of promoters defined as Tail-dependent (TD; this work) and coactivator-redundant (CR) (Donczew et al., 2020a). **(E)** Boxplot showing SAGA dependence of the two classes of CR genes: TD (Tail dependent) and TI (Tail-independent). Data from (Donczew et al., 2020a)

Our Med16 depletion results contrast with a recent report showing that Med16 plays a largely negative role (Saleh et al., 2021). In this earlier study, ∼90% and 70% of affected genes were upregulated by rapid Med16 depletion and deletion, respectively. The prior study used yeast with a different genetic background but whether this accounts for the difference is unknown. Here, we observed Tail subunit depletion to cause a transcription increase at Tail-independent genes, but only in the Med2/15 double degron strain (**Fig 1A; B**). We considered that this result might be due to a problem with the spike-in normalization for this RNA-seq experiment. However, repeating the experiment showed that the response of this strain to Med2/15 depletion was reproducible. The only other instance where we observed widespread but modest increase in transcription of Tail-independent genes upon Tail inactivation was when cells were stressed by brief growth in 1M NaCl (see below). In any event, we find that the Tail has a positive function at nearly all affected genes under the conditions tested with these two specific exceptions. The positive function of the Tail agrees with our results shown below where Tail depletion causes a defect in PIC formation specifically at the Tail-dependent genes and no change at Tail-independent genes.

### MED Tail and SAGA cooperate in transcription of Tail-dependent genes

Our prior work identified two sets of genes that differ in the requirements for the coactivators TFIID and SAGA (Donczew et al., 2020a). Expression from ∼87% of expressed genes, termed TFIID-dependent genes, is sensitive to rapid TFIID depletion while the remaining genes, termed coactivator redundant (CR) are moderately sensitive to TFIID or SAGA depletion. Transcription from this latter gene set is strongly decreased only upon simultaneous depletion of TFIID and SAGA. Remarkably, 95% of Tail-dependent genes are in the CR gene class, with 42% of CR genes being Tail-dependent (**Fig 1D**). Thus, the CR genes can be split into two roughly equal classes, Tail-dependent and Tail-independent genes. Analysis of these two classes of CR genes showed that expression from Tail-dependent genes is more strongly dependent on SAGA compared with Tail-independent CR genes (**Fig 1E**). After inactivation of SAGA by simultaneous depletion of Spt3/Spt7 (Donczew et al., 2020a), transcription of Tail-dependent genes decreased ∼ 1.7-fold compared with ∼1.2 fold for CR Tail-independent genes. Conversely, the Tail-dependent set of CR genes are less dependent on TFIID as determined by rapid Taf1 depletion (1.3 vs 1.8-fold) (**Fig S2C**). Interestingly, the Tail-dependent CR genes are on average insensitive to Bdf1/2 depletion (1.0 vs 1.2-fold) and are depleted for Bdf1 binding compared with the Tail-independent CR genes, which explains our prior observation that Bdf1/2 contribute to transcription of only a subset of CR genes (Donczew and Hahn, 2021). Together, our results suggest that Tail function is linked with the roles of SAGA, TFIID and Bdf1/2 in transcription initiation.

To further explore the relationship between MED Tail and SAGA, we measured transcription changes due to rapid depletion of Tail + SAGA (Med15/Spt7-degrons), Tail + TFIID (Med15/Taf1-degrons) and SAGA + TFIID (Spt7/Taf13-degrons) and compared these effects to changes caused by depletion of individual coactivators and Tail (**Fig 2A**; **Table S1**). This analysis revealed a striking cooperation between Tail and SAGA for expression of Tail-dependent genes. First, depletion of either Med15 (Med15-degron) or SAGA (Spt3/7-degrons) reduced transcription at Tail-dependent genes by ∼1.7-fold with little average effect on Tail-independent genes. However, co depletion of Tail and SAGA (Med15/Spt7-degron) caused a strong defect in transcription of Tail-dependent genes (∼3.2-fold), showing that SAGA and Tail cooperate in transcription of this gene set. Second, we found earlier that simultaneous depletion of TFIID and SAGA strongly reduced transcription of all Pol II transcribed genes (Spt7/Taf13-degrons) (Donczew et al., 2020a). Strikingly, we found a very similar transcription defect (∼7-fold) upon simultaneous depletion of Tail and TFIID (Med15/Taf1-degrons). This shows that, in the absence of TFIID, expression of the Tail-dependent genes strongly requires both Tail and SAGA. However, depletion of either Tail or SAGA in the presence of TFIID leads to a milder transcription defect, suggesting that TFIID can cooperate with Tail and SAGA for transcription of this gene set.

**Fig 2.**
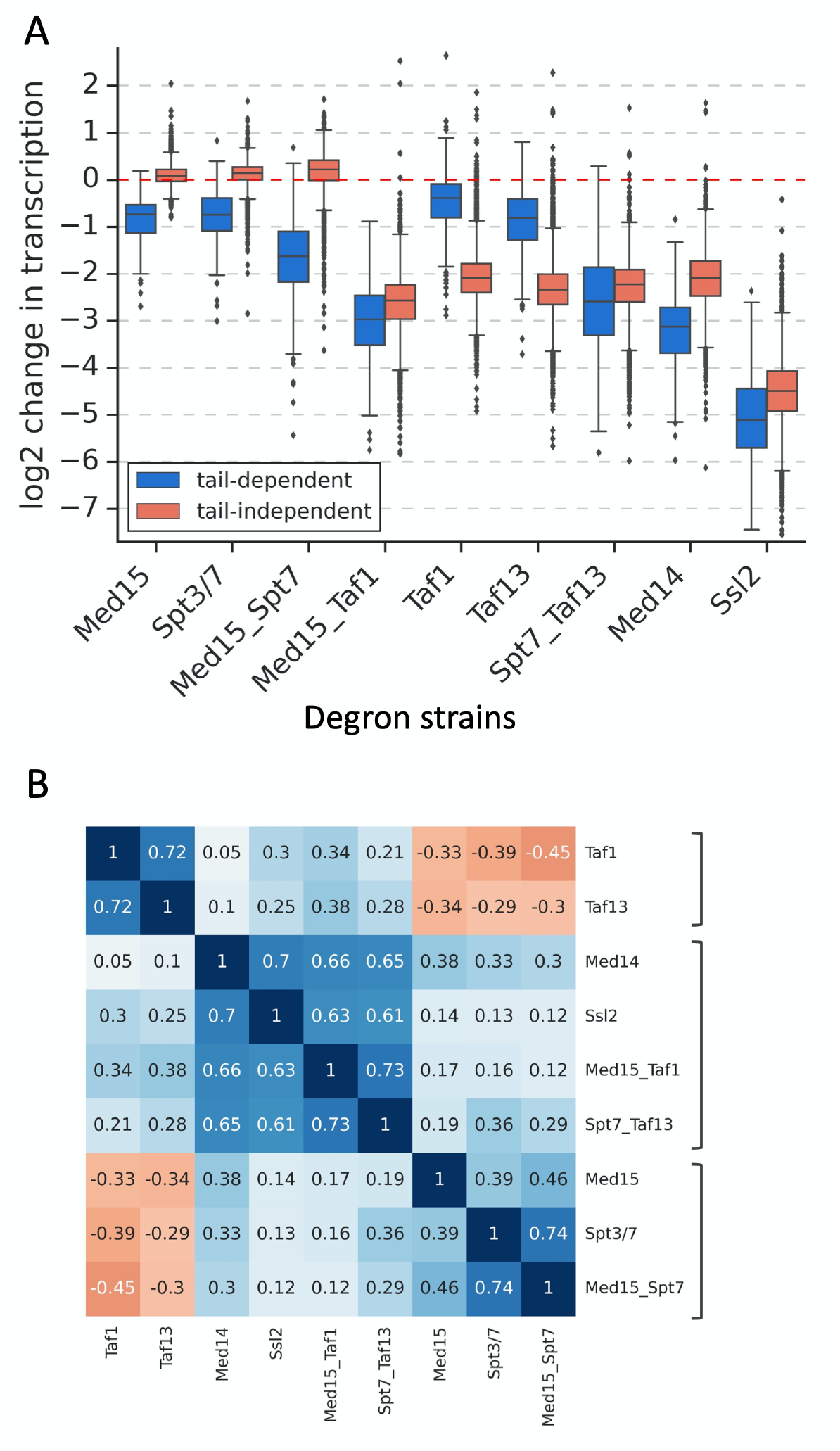
Cooperativity of Tail, SAGA and TFIID in transcription of Tail-dependent genes. **(A)** Boxplot showing log2 changes in transcription of Tail-dependent and Tail-independent genes after degron depletion of the indicated MED, SAGA and/or TFIID subunits. **(B)** Matrix of Spearman correlation coefficient values for log2 changes in transcription in the indicated experiments. Brackets indicate correlated groups.

The functional relationship between Tail, SAGA and TFIID is illustrated by the Spearman correlation matrix of transcriptional changes mediated by depletion of these and other factors (**Fig 2B**). Changes due to depletion of SAGA+TFIID and Tail + TFIID are in the same cluster as strong genome-wide transcription decreases caused by depletion of the basal factor TFIIH (Ssl2) and Med core (Med14). A separate cluster of changes due to Tail, SAGA or Tail + SAGA depletion highlights the linkage of Tail and SAGA function. In contrast, transcriptional changes caused by TFIID depletion cluster separately. Together, our results show that SAGA and Tail functions are linked and that TFIID, SAGA and Tail all contribute to transcription of the Tail-dependent genes.

### The set of Tail-dependent genes shows little change under stress conditions

While it’s surprising that there are only a small number of Tail-dependent genes, it’s possible that the composition of this gene set changes depending on which transcription factors are active under different growth conditions. For example, changes in levels or function of Tail-interacting transcription factors during stress response may lead to changes in the set of Tail-dependent genes. To test this model, we first subjected Med15-degron containing cells to three well-characterized stress conditions: a) heat shock at 37 deg. for 30 min (Morano et al., 2012). b) osmotic shock by addition of 1M NaCl for 30 min (Saito and Posas, 2012), or c) amino acid starvation using 0.5 μg/ml sulfometuron methyl (SM) for 60 min (Herbig et al., 2010; Jia et al., 2000; Natarajan et al., 1999). After these treatments, IAA was added for 30 min to deplete Med15 followed by 4-ThioU RNA-seq.

Without Med15 depletion, all three stress conditions caused modest global changes in transcription as well as induction or repression of stress-specific genes (**Table S1; Fig S3**). To characterize Tail-dependence in these new growth conditions, we confined our analysis to the set of 4885 genes for which mRNAs could be quantitated under non stressed growth as nearly all stress-induced genes were included in this gene set. Comparing results from all growth conditions, we surprisingly found little change in the set of Tail-dependent genes (**Fig 3A, B**). We found a clear separation in the behavior of the Tail-dependent and independent genes in response to Med15 depletion and that the majority of Tail-dependent genes show similar transcriptional changes with or without applied stress following Med15 depletion (**Fig 3C**). Although we have tested only a small number of all possible stresses, our results suggest that, for most genes, Tail-dependence is an inherent property of the gene and that only a small subset of mRNA genes are Tail-dependent.

**Fig 3.**
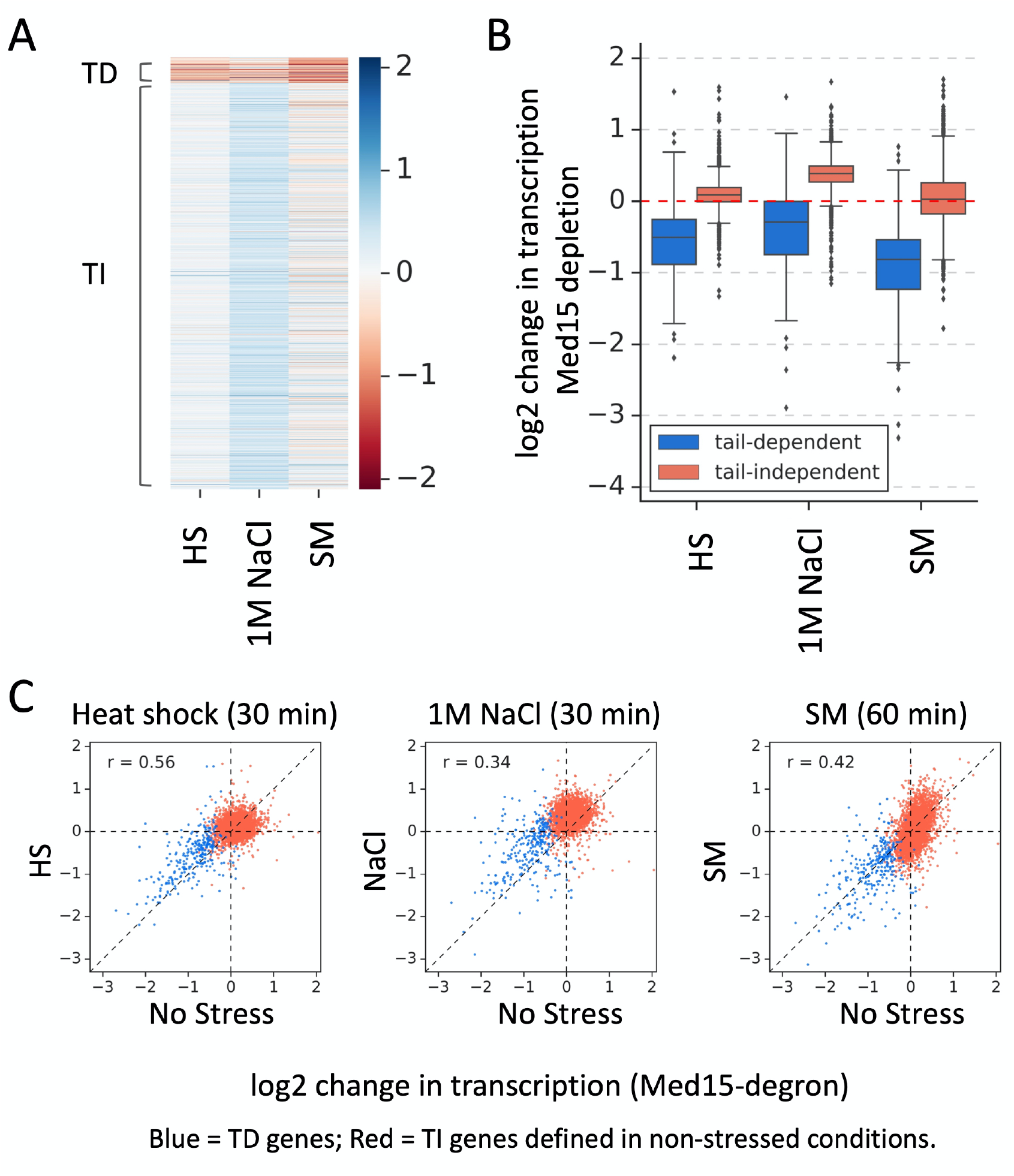
Little change in the identity of Tail-dependent genes during stress response. **(A)** Heatmap representation of Tail-dependent transcription under three stress conditions. Cells were briefly treated with the indicated stress conditions as described in the text and Methods followed by degron depletion of Med15. Transcription changes were assayed by 4-ThioU RNA-seq. Tail-dependent (TD) and Tail-independent (TI) genes from Fig 1 are marked. **(B)** Boxplot quantitating transcriptional changes after stress treatment. **(C)** Scatter plots comparing the response to Med15 depletion under indicated stress (Y-axis) vs no stress (X-axis) conditions. Points are labeled according to the Tail-dependent (blue) and Tail-independent (red) gene classes. Spearman’s rank correlation coefficients (r) for Tail-dependent genes are shown.

### Properties of Tail-dependent and independent genes

For insight into properties that dictate Tail dependence, we examined gene regulatory regions for features that are biased at Tail dependent or independent genes (**Table S2**). First, we found that the binding of MED to UAS elements, as determined using ChEC-seq with Med8 and Med15-MNase fusions, is enriched at Tail-dependent genes (**Fig S4A**). Out of 5891 genes analyzed for Med8 and Med15 binding by ChEC-seq (Donczew et al., 2020a), we found 1483 and 1337 UASs bound by Med8 and Med15, respectively. MED was found at the UAS from ∼60% of Tail-dependent genes, but at only ∼20% of Tail-independent genes. Further, of all the genes with UAS-bound MED, the Tail-dependent genes have on average 3.5-fold higher occupancy compared with Tail-independent genes (**Fig S4B**).

We found that the chromatin remodeler Swi/Snf was moderately enriched at Tail-dependent genes (**Fig S4C**). Conversely, the H4 acetylase NuA4 and its acetylation target H4 K12-Ac are enriched at Tail-independent genes (**Fig S4D, F**). These latter results are consistent with earlier findings that NuA4 and the H4-Ac mark are biased toward TFIID-dependent genes (Donczew and Hahn, 2021). Similarly, the chromatin remodeler SWR1 is strongly enriched at Tail-independent genes (**Fig S4E**). Finally, analysis of nucleosome and transcription factor binding data showed two properties that are distinct at Tail-dependent and independent genes (**Fig S4G, H**). First, the width of the nucleosome depleted region (NDR) in gene regulatory regions (Chereji et al., 2018) is widest at Tail-dependent genes (median distance 233 vs 132 bp). Likely related to this, we found that the UAS-TSS (transcription start site) distance (Rossi et al., 2021) was the longest at the Tail-dependent genes (median 179 vs 125 bp). Taken together, while we found no single feature or factor that is exclusive to Tail-dependent or independent genes, there are many features that are clearly biased at each gene class and at least some of these may contribute to Tail-dependence.

### Depletion of Middle subunit Med7 largely mimics Tail inactivation

To gain additional insight into Tail function we systematically depleted additional MED subunits from the Head and Middle modules to test their roles in genome-wide transcription. Analysis of 4-ThioU RNA-seq results showed that MED subunits can be clustered into four groups based on genome-wide expression changes (**Fig 4A, B; Fig S5A; Table S1**). Group 1 identifies MED subunits most important for transcription at both gene classes. The most strongly required subunits in this group are Med17 (Head module), Med10 (Middle module – a component of the Hook domain that binds CDK7) and, as shown earlier, the connector subunit Med14 (El_Khattabi et al., 2019; Petrenko et al., 2017; Warfield et al., 2017). Depletion of these 3 subunits led to a ≥4-fold decrease in genome-wide transcription. Depletion of two other subunits in group 1 caused a ∼2-fold transcription decrease: Med31 (Middle module - a component of the CTD-binding Knob domain and Med19 (Hook domain). For comparison, transcription decreases >4-fold upon depletion of the TFIIH subunit cyclin Ccl1 (**Fig 4A**) and decreases of >16-fold are observed upon depletion of TFIIA subunit Toa1 or the TFIIH translocase subunit Ssl2 (Donczew and Hahn, 2021). Since the extent of depletion for all these proteins is comparable, our results suggest that ≥80% of Pol II transcription is MED-dependent. The Tail-dependent genes are on average ∼1.5-fold more sensitive to depletion of these MED subunits and basal factors compared with Tail-independent genes (**Fig 4A**). The only exception is for Med19 where Tail-independent genes are slightly more sensitive to Med19 depletion.

**Fig 4.**
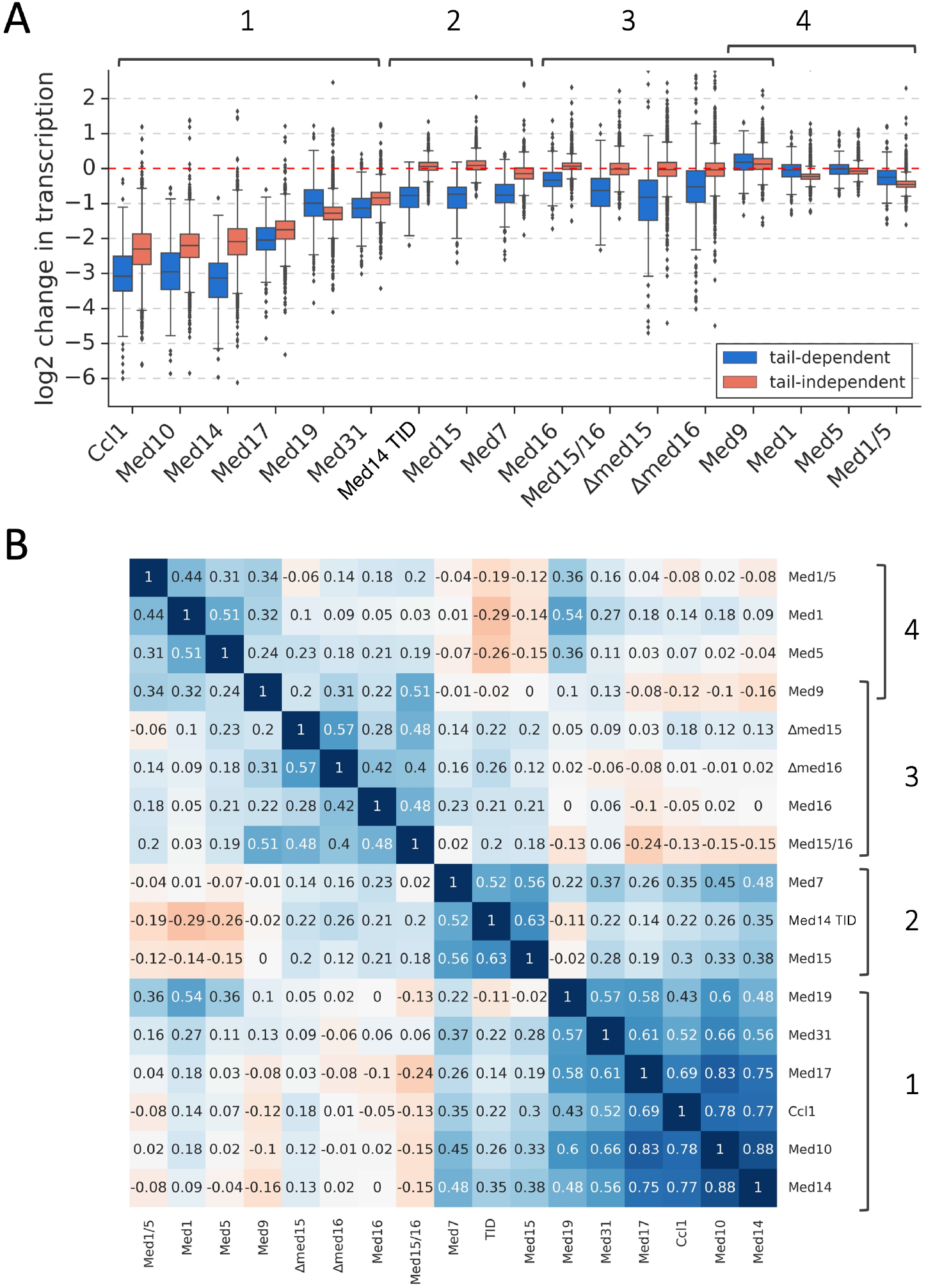
Mediator subunits can be divided into four groups based on the changes to genome- wide transcription following rapid depletion. **(A)** Boxplot showing log2 change in transcription for 4885 genes measured by 4-ThioU RNA-seq after degron-depletion or deletion of indicated factors. Genes are grouped into Tail-dependent and Tail-independent categories. Four distinct clusters of MED subunits defined in this work are marked. **(B)** Matrix of Spearman correlation coefficient values between log2 changes in transcription in indicated experiments. Four distinct clusters of MED subunits defined in this work are marked.

A recent report suggested that rapid Med14 depletion had a smaller effect on genome-wide expression compared with disruption of the Head module by Med17 depletion (Tourigny et al., 2021). This result was used to suggest that the Head module retained partial function in the absence of intact MED core. In contrast, we find near-equivalent effects of Med14 and Med17 depletion on Tail-independent genes and that Tail-dependent genes are slightly more sensitive to Med14 depletion compared with Med17 depletion. The reason for this discrepancy is unknown, however, our results are consistent with prior biochemical assays showing that Head function requires both the Middle module and Med14 (Cevher et al., 2014; Plaschka et al., 2015).

The remaining three MED subunit groups all contain Tail subunits. Depletion of the group 2 subunits Med15, the Med14 TID and the Middle subunit Med7 specifically decrease transcription of the Tail-dependent genes and the effects of subunit depletion within this set are well correlated. The inclusion of Med7 in this group (r= 0.52-0.56) was surprising as Med7 is an essential subunit that is physically distant from Tail (**Fig S5A**). Med7 is an extended polypeptide that can be separated into two regions (Baumli et al., 2005; Koschubs et al., 2009; Nozawa et al., 2017). The Med7 N-terminal domain, together with Med31, forms the Knob domain. The C-terminal Med7 region can be divided into two helical domains separated by a flexible hinge. The first helical domain, together with Med21, forms a 4-helix bundle that comprise the upper half of the Hook while the C-terminal helix, together with Med21, forms an extended coiled-coil that connects to Med4 and Med9 in the Plank domain. Mutations in the Med7 hinge region cause changes in conformational flexibility of MED and reduce the affinity for Pol II (Sato et al., 2016). Based on our results, it seems likely that depletion of Med7 interferes with functions related to the role of MED Tail. Med21 and Med4 are also components of the long helical region of the Middle module. However, we were unable to test whether depletion of these subunits had similar defects as Med7 since we could not obtain strains with C-terminal degron fusions to these subunits.

The third group of MED subunits includes the Med16-degron and the Med15 and Med16 deletions. As noted above, expression changes in the Med16 degron and deletion strains modestly correlate, while the Med15 degron and deletion strains show only weak correlation. Med16 was originally identified using a reporter assay where deletion of Med16 both positively and negatively affected expression (Jiang and Stillman, 1992; Sternberg et al., 1987). Elimination of Med16 has also been found to weaken the connection between the MED Tail and the MED core domain (Béve et al., 2005; Cevher et al., 2014; Li et al., 1995; Zhang et al., 2004). Given this, it was surprising that rapid depletion of Med16 resulted in only weak decreases in transcription from the Tail-dependent genes while most Tail-independent genes showed no significant change in expression. Similar genome-wide averages were observed in the Med16 deletion strain.

The fourth group contains subunits Med1, Med5 and Med9, with the latter subunit showing modest to weak correlations in expression changes with both group 3 and 4 subunits (r = 0.5-0.2). Med5 is a more peripheral Tail subunit that connects the Tail with Med1, which in turn interacts with Middle subunits Med4/9 in the Plank (**Fig S5A**). Expression changes after Med5 and Med9 depletion are minimal and transcription from Tail-independent genes is only weakly decreased after Med1 depletion. Simultaneous depletion of Med1 and Med5 gives weak but slightly stronger transcription changes than observed in the single degron strains. The lack of strong expression changes upon Med1 depletion was surprising as Med1 is a common activator target in higher eukaryotes and earlier results from mammalian cells showed that transcription from a subset of genes is affected by Med1 depletion (El_Khattabi et al., 2019).

To examine how Med7 contributes to Tail function, we asked which regions of Med7 are specifically important for transcription of Tail-dependent genes. It was shown earlier that simultaneous expression of separate N and C-terminal segments of Med7 can reconstitute function (Koschubs et al., 2009). Based on this finding, we expressed Med7 residues 1-102 (Med7N) and 98-222 (Med7C) on separate plasmids, with both under control of the Med7 promoter and in strains where one of these regions is fused to a degron. Upon depletion of Med7N, which is situated in the Knob domain, we found that transcription of both Tail-dependent and independent genes was decreased ∼2-fold (**Fig S5B**). In contrast, depletion of Med7C led to greater defects in expression from Tail-dependent vs Tail-independent genes (∼2-fold vs 1.4-fold). From this, we conclude that the Tail-specific defect resulting from Med7 depletion is mostly mediated by depletion of the C-terminal region.

To explore the consequences of Med7 depletion on MED integrity, we used IP assays to monitor association of subunits after depletion of Med14 N (the domain connecting Head and Middle), Med14 TID, or Med7. From structural considerations and prior studies, Med14 N should be required for integrity of the entire complex, while TID depletion is expected to cause Tail dissociation. Strains containing degrons on either the Med14 N terminal domain (Med14N; residues 1-746), the Med14 TID or Med7, in conjunction with epitope tags on selected Med subunits were treated with either DMSO or IAA. Extracts were generated and immunoprecipitated using polyclonal antisera specific for Med17 (Head) or Med3 (Tail) and then probed for co precipitation of Tail (Med2,15, 16), Middle (Med1, 10 and/or 7) and Head (Med18). Results are summarized in **Fig 5A** with data shown in **Fig S6**. As expected, depletion of Med14N caused dissociation of Head, Middle and Tail while leaving Tail integrity intact. For example, after depletion of Med14N, Med17 was efficiently precipitated with Med18 (Head) but association was strongly reduced with Med10 (Middle) and all tested Tail subunits. Additionally, after Med14 N depletion, Med3 still strongly associated with Tail subunits Med2, 15, and 16 showing that Tail remained intact. In contrast, depletion of Med14 TID resulted in dissociation of Tail from MED core. For example, we found that interaction of Head and Middle was unaffected by TID depletion as Med17 co precipitated with Med18, 1, and 10. In contrast, Head association was strongly reduced with all tested Tail subunits (Med2, 15, 16). In addition, the Tail was partly disrupted by TID depletion as Med3-Med16 association was strongly reduced. Finally, depletion of Med7 left MED largely intact as Head, Middle and Tail subunits all co precipitated. Except for Med1 and Med10, that showed reduced association after Med7 depletion, all tested Head and Tail subunits tested co precipitated with Med17 and Med3. These results, combined with our earlier finding that Med7 depletion causes a defect in transcription of Tail-dependent genes, indicate that Med7 depletion causes a defect in MED Tail function that is not mediated by the physical loss of the Tail module.

**Fig 5.**
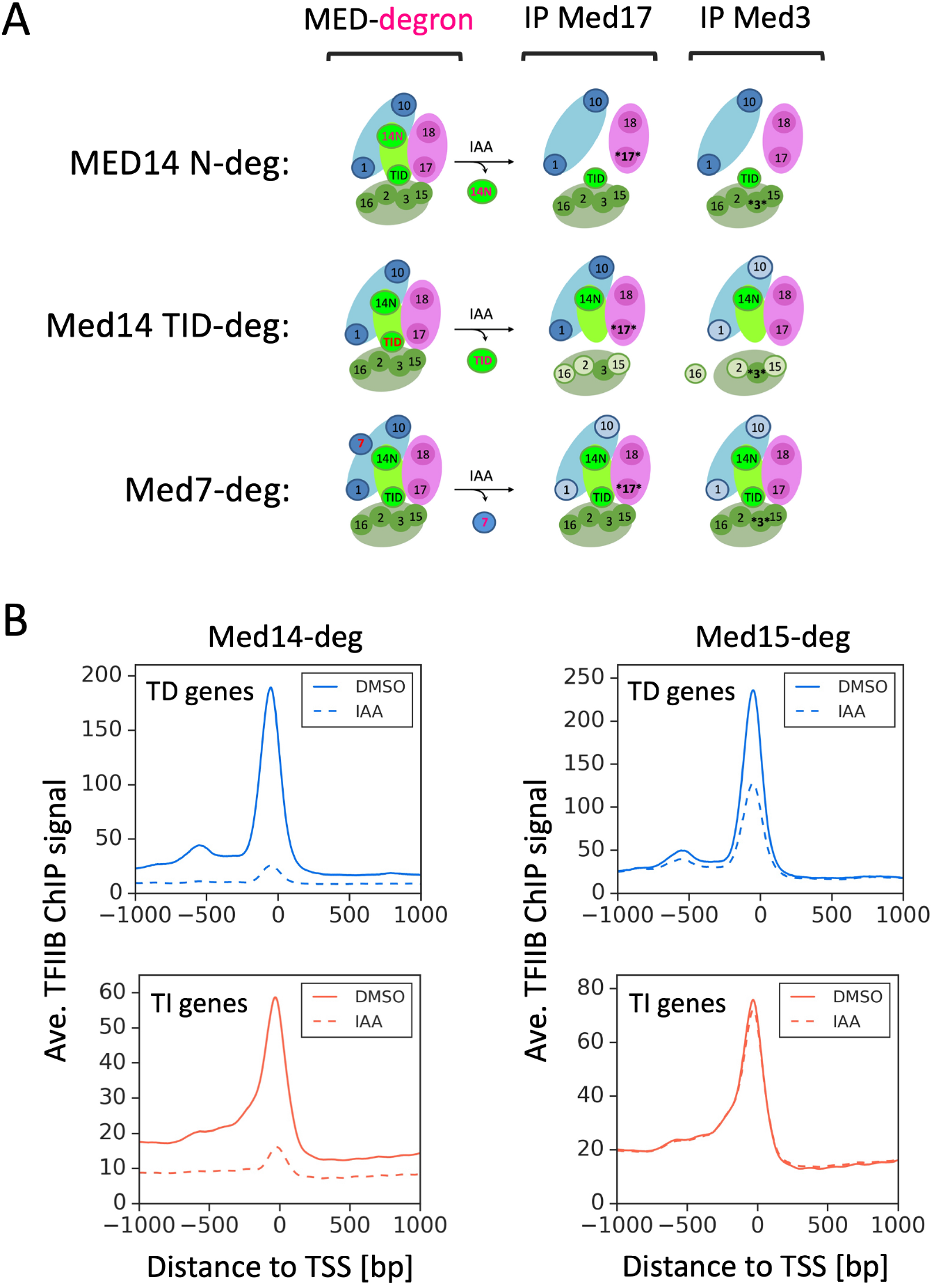
MED integrity after subunit depletion and the role of MED core and Tail in PIC formation. **(A)** Schematic summarizing dissociation of Mediator modules after degron-depletion of either Med14N, Med14 TID or Med7. IPs of either Med17 (Head) or Med3 (Tail) were performed with extracts prepared from strains depleted of either Med14N, Med14 TID or Med7. Mediator subunits that co precipitated were determined by probing for Flag-tagged Med2, 15, 16 (Tail), Med1, 10 (Middle) and Med18 (Head). Figure is based on IP data from **Fig S6. (B)** Average plots comparing TFIIB ChIP-seq signals before (DMSO solid line) and after (IAA, dashed line) Med14 or Med15 depletion at 4885 genes classified into Tail-dependent (TD) and Tail-independent (TI) categories.

Finally, we assayed expression changes in strains with MED mutations that have previously been linked to transcriptional regulation. The EWE (Expression Without heat shock Element) mutations were identified as allowing expression of a reporter gene with a partially defective UAS (Kremer et al., 2012; Singh et al., 2006). EWE phenotypes were caused by mutations in Med7, Med21, Med10, Med14 and Med19 and by removal of the Med14 TID via nonsense mutation. We found that the *med19* EWE mutant had a very weak phenotype and the *med10* EWE mutation was lethal in our strain background but the EWE mutations in *med7* and *med21* showed slow growth and temperature sensitive phenotypes as reported earlier (**Fig S1A**). Analysis of these latter two mutants by 4-ThioU RNA-seq did not reveal gene-specific transcription defects as the mRNA levels of most genes were decreased >2-fold (**Fig S5B**). Finally, multiple missense mutations within the Med6 N-terminal *α* helices were reported to produce a defect in activated transcription (Lee et al., 1997). This region of human Med6 was recently shown to interact with and assist in positioning the TFIIH kinase module (Abdella et al., 2021; Chen et al., 2021; Rengachari et al., 2021). Deletion of the first 2-4 helical regions of Med6 (residues 24-33; 39-51; 54-107), the sites of the multiple mutations comprising the *med6* ts alleles used in the earlier study (Larivière et al., 2012), was lethal in our strain background. However, we found that a shorter deletion (residues 13-20) that eliminated only the first *α* helix was viable but showed moderately slow growth and was cold sensitive. RNA-seq analysis found that this latter mutation caused a general defect in transcription rather than gene-specific effects (**Fig S5B**). In summary, the only mutation or subunit depletion we found outside of the Tail module that results in strong gene-specific transcription defects was depletion of the Middle subunit Med7.

### Tail stimulates PIC formation at Tail-dependent genes

Some prior studies suggested that strains with Tail subunit deletions have modest decreases in PIC formation at most genes and a broad genome-wide decrease in Pol II ChIP signals (Jeronimo et al., 2016; Knoll et al., 2018). To examine how rapid Tail subunit depletion affects PIC formation we depleted Med14 or Med15 and used ChIP-seq to probe for TFIIB-DNA binding as a proxy for monitoring defects in PIC formation (**Fig 5B; Table S3**). We found that Med14 depletion strongly decreased TFIIB binding at all active genes while depletion of Med15 reduced TFIIB binding (∼1.8-fold) only at Tail-dependent genes. These results show that MED function is critical for PIC formation at all genes, while the Tail has a moderate contribution to PIC formation only at the Tail-dependent genes – in excellent agreement with the effects of rapid MED subunit depletion on transcription shown above.

### Med10, Med19-MNase probes detect MED at promoters of most active genes

As noted above, prior ChIP and ChEC assays found that yeast MED maps to UAS elements rather than core promoters, unless CDK7/Kin28 is inhibited or depleted. This leaves open the question of how widespread direct association of MED with promoters is under normal growth conditions. The structures of yeast and human MED-PIC complexes indicate that MED is positioned far from promoter DNA (Abdella et al., 2021; Chen et al., 2021; Rengachari et al., 2021; Robinson et al., 2016; Schilbach et al., 2017) and this, along with the short PIC lifetime in vivo (Nguyen et al., 2021), may explain difficulties in detecting MED at promoters. To attempt detection of MED-promoter binding, MNase fusions were made with Hook domain subunits Med10 and Med19. In the PIC, these two subunits are positioned ∼140 Å distant from downstream promoter DNA but this DNA is predicted to be readily accessible for DNA cleavage. In agreement with the findings that MED is important for transcription of all mRNAs, both these MNase fusions cleaved DNA at many promoters under standard growth conditions. DNA cleavage overlapped with TFIIB ChIP-seq signals at many genes and at locations distinct from all other Med-MNase fusions tested (**Fig 6A; Fig S7A; Table S3**). As expected, Med10-MNase cleavage was reduced to near background levels upon depletion of Med14. Using our prior quantitative criteria for MED binding, we observed MED at ∼60% of all mRNA coding genes (3156 promoters) with no bias toward Tail-dependent or independent genes. This contrasts with the Tail-dependent bias observed for other Med-MNase fusions that cleave DNA at the UAS (**Fig 6B; S4A**).

**Fig 6.**
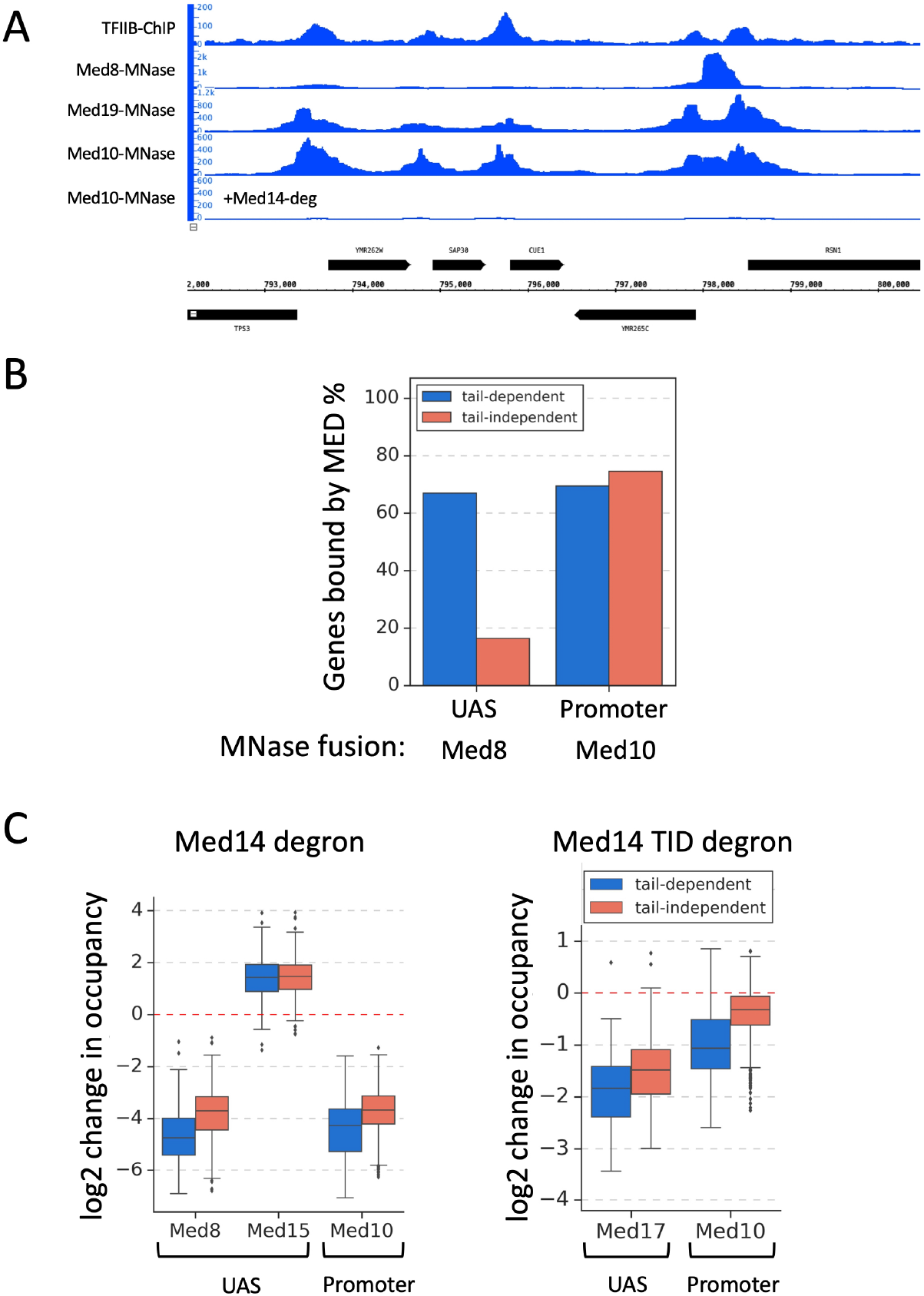
Assay for MED-promoter binding and the roles of MED core and Tail in MED-UAS and promoter binding. **(A)** Genome browser image showing comparison of TFIIB ChIP-seq and Med8, Med10, and Med19 ChEC-seq signals at a representative genomic location. Also shown are Med10-MNase cleavage signals after degron-depletion of Med14. **(B)** Bar plot showing percentage of UASs and core promoters in each gene class (Tail-dependent and Tail-independent) bound by Mediator. Med8 signal represents Mediator occupancy at UAS and Med10 signal at core promoter. **(C)** MED binding was monitored to UAS and promoters using the indicated Med-MNase strains. Results are shown as log2 change in MED ChEC-seq signals (interpreted as occupancy) after depletion of either Med14 or the Med14 TID. Data from both Tail-dependent and Tail-independent genes are shown.

### MED is recruited to most promoters via a Tail independent pathway

Using this new assay, we investigated the relative importance of Tail for recruitment of MED to core promoters. We first examined the relationship between transcription output and MED binding at promoters and UASs. For this analysis, we confined our comparisons to actively transcribed genes which had an associated MED peak based on Med10 (promoter) or Med8 (UAS) ChEC-seq results (173 promoters and 186 UASs at Tail-dependent genes; 2681 promoters and 865 UASs at Tail-independent genes). This latter set encompasses UASs from only the ∼20% of Tail-independent genes where MED binding was observed. For comparison, we found good correlations between transcription levels and signals from Pol II ChIP (r = 0.66) and TFIIB ChIP (r = 0.58) (**Fig S7B, C; Table S3**).

**Figure S7D, F** shows that there is a modest correlation between transcription levels and the occupancy of MED bound to Tail-dependent promoters (r = 0.43) with a weaker correlation at promoters of Tail-independent genes (r = 0.21). A difference between the two gene classes was also observed for MED binding to UASs. There was a weak correlation between transcription output and the level of MED-UAS binding at Tail-dependent genes, but no correlation at Tail-independent genes. This suggests that occupancy of MED binding at promoters and UASs is partially limiting for transcription of Tail-dependent genes. In contrast, the level of MED-promoter binding seems only partially limiting for transcription of Tail-independent genes and MED-UAS binding is not limiting at these genes.

To further explore the relationship between Tail and MED binding, we used ChEC-seq to monitor MED promoter and UAS binding after Tail (Med14 TID) depletion (**Fig 6C; Table S3**). As a control, we found that Med14 depletion caused a severe reduction in MED core binding to both UASs and promoters but that Tail (Med15) binding to UASs was mildly increased. This latter finding agrees with earlier results showing Med15-UAS binding in the absence of MED core (Yarrington et al., 2020; Zhang et al., 2004). The effects we observed were nearly equivalent at both the Tail-dependent and independent genes. In contrast, depletion of Tail function by the TID degron caused different defects for UAS and promoter binding. UAS binding of both gene classes was reduced 2-4 fold upon TID depletion. However, MED-promoter binding was most affected at the Tail-dependent genes. Monitored by Med10-ChEC assays, promoter binding was reduced ∼2-fold at Tail-dependent genes upon TID depletion but only by ∼1.3-fold at Tail independent genes. These binding changes are consistent with the effects of TID depletion on levels of newly synthesized mRNAs and PIC formation where transcription and TFIIB recruitment are insensitive to TID depletion at Tail-independent genes (**Fig 1B; 5B**). Together, our results show that MED can be recruited to most Tail-independent promoters in the absence of Tail function and UAS binding while MED-UAS binding is important for transcription of only Tail-dependent genes.

## Discussion

The role of MED in transcription activation of a small number of exemplary genes led to the general consensus that one of the most important functions of MED is a direct role in transcription activation. (Allen and Taatjes, 2015; Jeronimo and Robert, 2017; Soutourina, 2018). Consistent with the model that gene activation involves activator-MED binding, MED contains activator-binding subunits, and these are important for the transcriptional response to specific activators. In opposition to the view that this is a broadly used mechanism, most MED activator-binding subunits are not essential for viability. While the reported transcriptional changes upon depletion of these subunits have been variable, expression from only a subset of genes is affected by elimination of these activator-binding subunits. Here, we have investigated the genome-wide role of MED in gene regulation by rapidly disrupting the yeast MED activator binding domain (the Tail module) and observing defects in genome-wide nascent transcription. This, coupled with a new assay that readily detects MED at promoters, revealed that only a small percentage of yeast genes follow the consensus model where activator-mediated MED recruitment to UAS elements is important for transcription. Rather, at most genes, MED bypasses the UAS and associates with promoters where it plays key roles in PIC formation and initiation rather than a direct role in transcription activation. Tail-independent recruitment of MED to promoters had been previously observed upon inhibition of transcription initiation (Jeronimo et al., 2016; Knoll et al., 2018; Petrenko et al., 2016) but it was proposed that this was an alternative to the dominant pathway via UAS recruitment. Our findings suggest that Tail-independent recruitment of MED is the predominant pathway used at nearly all genes and they have broad implications for the roles of MED and mechanisms of transcriptional regulation.

A striking finding from our studies is that expression from only ∼6% of yeast genes is sensitive to Tail disruption. This was surprising given that it has been estimated that ∼40-50% of yeast transcription factors contain acidic activation domains that typically bind Med15, the principal activator-binding subunit of yeast MED (Erijman et al., 2020; Sanborn et al., 2021). It was recently proposed, based on formaldehyde crosslinking experiments, that yeast sequence specific transcription factors regulate only ∼1,000 genes (termed STM genes) while the remaining 80% of genes were either bound by factors that function to exclude nucleosomes or that were devoid of detectable factors (Rossi et al., 2021). Even if this model is correct, the set of Tail-dependent genes (n =287) is much smaller than the set of STM genes. Unexpectedly, we found that the set of Tail-dependent genes changed little after yeast were exposed to several stress conditions that activated different transcriptional programs. This suggests that, for most genes, Tail-dependence is an inherent property of the gene regulatory region. We speculate that Tail dependence may be mediated by a combination of regulatory transcription factors, cofactor utilization, promoter sequence and chromatin configuration.

Our combined results strongly suggest that MED bypasses UAS binding at most Tail-independent genes and is directly recruited to promoters. First, MED binding to UASs is clearly biased toward Tail-dependent genes, with ∼60% of these genes having detectable MED at their UASs compared to only ∼20% of Tail-independent genes. MED-UAS occupancy is also much higher at Tail-dependent vs independent genes. In contrast, using MNase fusions to subunits in the MED Hook domain, we observe MED-promoter binding at most active genes without bias for either gene class. Second, at the ∼20% of Tail-independent genes that have detectable MED-UAS binding, we find no correlation between binding and transcription level, unlike the situation at Tail-dependent genes. Third, rapid depletion of Tail causes a strong decrease in MED-UAS binding at all genes but only a strong decrease in MED-promoter binding at Tail-dependent genes. This agrees with the finding that Tail disruption causes little or no change in transcription and PIC formation at Tail-independent genes. Coupled with the result that MED core is critical for PIC formation at all genes, our results suggest that the principal role of MED at most genes is in general transcription initiation rather than direct involvement in transcription activation. Based on prior biochemical findings and results shown here, the most important roles of MED are likely stabilization of the PIC and positioning/activation of the CTD kinase CDK7. Since much of MED seems associated with soluble Pol II and the affinity of MED for Pol II is strong, we imagine that Pol II and MED are often co recruited during PIC formation. At most Tail-independent genes, rather than targeting MED recruitment, we speculate that a critical function of sequence-specific transcription factors is to recruit the H4 HAT complex NuA4 that primarily targets acetylation of the +1 and -1 nucleosomes. We showed earlier that Bdf1/2, analogous to mammalian BET factors that bind acetylated H4, are critical for transcription of these genes and that H4 acetylation is important (but not absolutely required) for Bdf recruitment (Donczew and Hahn, 2021). However, at the Tail-dependent genes, our results are consistent with prior models for the direct role of MED in transcription activation. For example, dynamic binding of activators to MED Tail likely recruits a cluster of MED that assists in PIC formation and transcription initiation at these genes.

Our work also uncovered striking relationships between the functions of MED Tail, SAGA, TFIID and Bdf1/2 and suggest that there are three major classes of yeast Pol II-transcribed genes that are distinguished by their PIC assembly pathways and the roles of coactivators and BET factors (**Fig 7**). TFIID-dependent genes are the largest gene category (∼87% of Pol II genes). Transcription of these genes is insensitive to rapid SAGA and Tail depletion but strongly dependent on TFIID. The CR gene class that utilizes both SAGA and TFIID (∼13% of genes) can be split into two near equal classes: Tail-independent and Tail-dependent. MED directly binds promoters of the Tail-independent CR genes, and these genes are more dependent on TFIID and less dependent on SAGA compared to the Tail-dependent CR genes. The Tail-independent CR genes also have a modest dependence on Bdf1/2 while the Tail-dependent genes are insensitive to Bdf depletion. The Tail-dependent genes are distinguished by their dependence on MED Tail, MED-UAS binding and stronger dependence on SAGA but weaker dependence on TFIID. SAGA and Tail cooperate in transcription of these genes as simultaneous depletion of Tail and SAGA have a greater effect than depletion of either factor alone. Additionally, co-depletion of SAGA + Tail, SAGA + TFIID and Tail + TFIID, uncovered an unexpected role for TFIID at these genes: when TFIID is absent, there is a very strong requirement for both Tail and SAGA. This suggests that TFIID, SAGA and MED Tail can cooperate under conditions where one of the other cofactors is absent. Combined, these gene classes set the parameters for the genome-wide roles of transcription factors, coactivators, and H4 acetylation at different subsets of genes.

**Fig 7.**
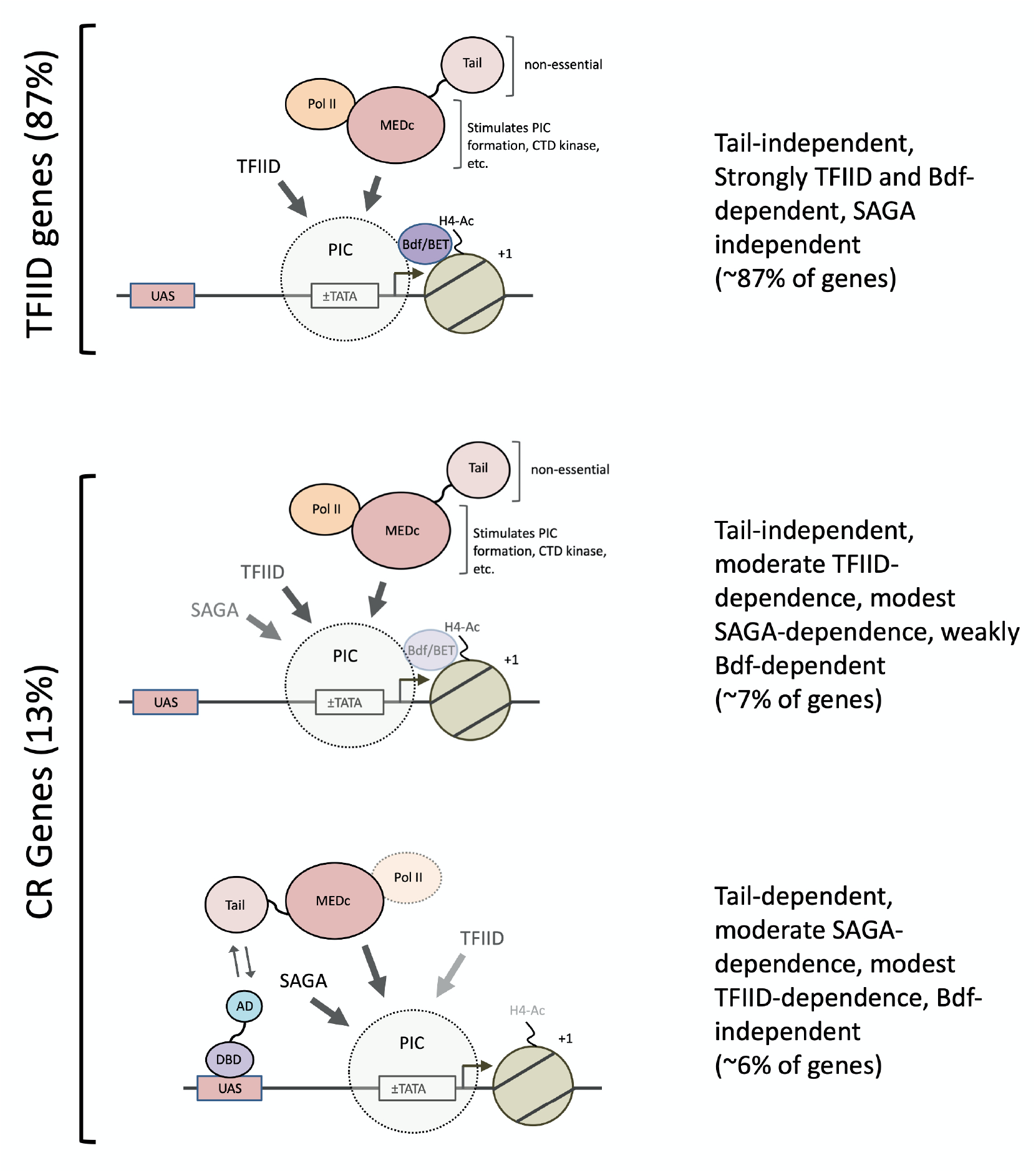
**Three classes of yeast genes, their coactivator requirements and MED recruitment pathways**. Genes (n = 4885) are separated into TFIID-dependent, CR Tail-independent, and CR Tail-dependent classes. The relative dependence of transcription on TFIID, SAGA, Bdf1/2 and H4 acetylation is indicated by transparency. Coactivator and Bdf dependence are defined here as transcription changes after a 30 min depletion (Donczew et al., 2020a)(Donczew and Hahn, 2021) Long-term depletion of SAGA reduces transcription from all mRNA genes due to changes in chromatin modifications.

Finally, our analysis of MED subunit function in the Head and Middle modules revealed that the Middle module subunit Med7 plays an important role in Tail function. We found that rapid depletion of Med7 gave a pattern of gene-specific transcription defects very similar to Tail disruption and that this function is largely localized to the Med7 C-terminal helical region. This Med7-Tail connection was surprising as Med7 is located far from the Tail module. Med7 is an essential subunit and, from the MED structure, it should be required for assembly of the Middle module. We found that rapid Med7 depletion leaves MED largely intact but speculate that its long-term absence would inhibit assembly of nascent MED leading to lethality. It was earlier proposed that activator-MED binding may cause a conformational change in MED leading to a more active state. Importantly, it’s known that MED undergoes a conformational shift upon Pol II binding and that the Med7 hinge region, located in the C-terminal helical region is important for Pol II binding (Schilbach et al., 2017; Tsai et al., 2017). Our results with Med7 are consistent with the conformational change model, but further experiments to understand the role of Med7 in activator-mediated transcription and MED function will be an important subject for a future study.

## Methods

### Yeast strains and cell growth

Yeast strains are listed in **Table S4** and are all derivatives of BY4705 (Brachmann et al., 1998). Degron and MNase-tagged strains were constructed as previously described (Donczew et al., 2020a; Grünberg et al., 2016). However, in several instances, tagging proteins with the full-length IAA7 degron led to a growth phenotype. In these cases, proteins were tagged with a shortened IAA7 degron (termed mini degron in **Table S4**) that resulted in little or no growth phenotypes for the strains used in this work. The sequence of the mini degron tag is below. Small letters indicate the 3X V5 epitope tag, bold letters are linker sequence, and non-bold capital letters encode the shortened IAA7 degron:

ggtaaacctatacctaatccattattgggactagatggaaaaccaataccaaatcccttacttg gtttggattctacaccaattcctaatcctctattaggactggatagtaca**GGTGCCGGTGCTGG TGCCGGAGCTGGCGCAGGTGCT**AAGGAAAAATCTGCGTGTCCAAAGGACCCTGCAAAACCACCA GCCAAGGCACAAGTTGTAGGTTGGCCCCCTGTAAGATCCTATAGAAAGAATGTTATGGTTTCTTGCCAAAAATCTTCTGGAGGCCCTGAAGCAGCTGCATTTGTTAAAGTTAGTATGGACGGTGCTCCTTACTTGAGAAAAATAGACTTGAGAATGTATAAA

*S. cerevisiae* strains were grown as indicated in synthetic complete (SC) media (per liter: 1.7 g yeast nitrogen base without ammonium sulfate or amino acids (BD Difco), 5 g ammonium sulfate, 40 μg/ml adenine sulfate, 0.6 g amino acid dropout mix (without -Ile -Val) and supplemented with two micrograms/ml uracil and 0.01% other amino acids to complement auxotrophic markers). Standard amino acid dropout mix contains 2 g each of Tyr, Ser, Val, Ile, Phe, Asp, Pro and 4 g each of Arg and Thr. *S. pombe* strains were grown in YE media (0.5% yeast extract, 3% glucose). Where indicated, *S. cerevisiae* strains at an A600 of ∼ 1.0 were treated with 500 μM indole-3-acetic acid (IAA) dissolved in DMSO (or with DMSO alone) for 30 min. Where indicated, cells were exposed to stress conditions by incubation with 0.5 μg/ml sulfometuron methyl (SM) in DMSO (SM) or with DMSO alone for 60 min; 1 M NaCl for 30 min; or by addition of an equal volume of 44°C media to the 30°C culture followed by incubation at 37°C for 30 min as described in the text and figure legends, prior to RNA labeling.

### Western blot analysis

1 ml cell culture was collected and pelleted from strains after treatment with IAA or DMSO, incubated in 200 μl 0.1 M NaOH for 5 min at room temp, then resuspended in 100 μl yeast whole cell extract buffer (0.06 M Tris-HCl, pH 6.8, 10% glycerol, 2% SDS, 5% 2-mercaptoethanol, 0.0025% bromophenol blue). After heating for 5 min at 95°C, samples were centrifuged for 5 min at max speed, whole cell extracts were separated by SDS-PAGE and analyzed by Western blot using mouse monoclonal (*α*-Flag, *α*-V5) or rabbit polyclonal (*α*-Med17, *α*-Med3, *α*-Tfg2) antibodies. Protein signals were visualized by using the Odyssey CLx scanner and quantified using Odyssey Image Studio software (Li-Cor) by generating a standard curve using a titration from WT extract.

### RNA labeling and mRNA purification

All experiments were done in triplicate except the following samples which were done in duplicates: Med7_EWE3, med15_16_WT, *MED15* deletion, *MED16* deletion, Med21_EWE4, and Med21_WT. Newly synthesized RNAs were labeled as previously described (Bonnet et al., 2014). 10 ml *S. cerevisiae* or 20 ml *S. pombe* cells were labeled with 5 mM 4-Thiouracil (Sigma-Aldrich) for 5 min, the cells were pelleted at 3000 x g for 2 min, flash-frozen in liquid N_2_, and then stored at −80°C until further use. *S. cerevisiae* and *S. pombe* cells were mixed in an 8:1 ratio and total RNA was extracted using the RiboPure yeast kit (Ambion, Life Technologies) using the following volumes: 480 μl lysis buffer, 48 μl 10% SDS, 480 μl phenol:CHCl_3_:isoamyl alcohol (25:24:1) per *S. cerevisiae* pellet + 50 μl *S. pombe* (from a single *S. pombe* pellet resuspended in 850 μl lysis buffer). Cells were lysed using 1.25 ml zirconia/silica beads in a Mini Beadbeater-96 (BioSpec Products) for 3 min followed by 1 min rest on ice. This bead beating cycle was repeated twice for a total of 3 times. Lysates were spun for 5 min at 16K x g, then the following volumes combined in a 5 ml tube: 400 μl supernatant, 1400 μl binding buffer, 940 μl 100% ethanol. Samples were processed through Ambion filter cartridges until all sample was loaded, then washed with 700 μl Wash Solution 1, and twice with 500 μl Wash Solution 2/3. After a final spin to remove residual ethanol, RNA was eluted with 25 μl 95°C preheated Elution Solution. The elution step was repeated, and eluates combined. RNA was then treated with DNaseI using 6 μl DNaseI buffer and 4 μl DNaseI for 30 min at 37°C, then treated with Inactivation Reagent for 5 min at RT. RNA was then biotinylated essentially as described (Duffy and Simon, 2016; Duffy et al., 2015) using 40 μl (∼40 μg) total RNA and 4 μg MTSEA biotin-XX (Biotium) in the following reaction: 40 μl total 4-ThioU-labeled RNA, 20 mM HEPES, 1 mM EDTA, 4 μg MTSEA biotin-XX (80 μl 50 μg/ml diluted stock) in a 400 μl final volume. Biotinylation reactions occurred for 30 min at RT with rotation and under foil. Unreacted MTS-biotin was removed by phenol/CHCl_3_/isoamyl alcohol extraction. RNA was precipitated with isopropanol and resuspended in 100 μl nuclease-free H_2_O. Biotinylated RNA was purified also as described (Duffy and Simon, 2016) using 80 μl MyOne Streptavidin C1 Dynabeads (Invitrogen) + 100 μl biotinylated RNA for 15 min at RT with rotation and under foil. Prior to use, MyOne Streptavidin beads were washed in a single batch with 3 × 3 ml H_2_O, 3 × 3 ml High Salt Wash Buffer (100 mM Tris, 7.4, 10 mM EDTA, 1 M NaCl, 0.05% Tween-20), blocked in 4 ml High Salt Wash Buffer containing 40 ng/μl glycogen for 1 hr at RT, then resuspended to the original volume in High Salt Wash Buffer. After incubation with biotinylated RNA, the beads were washed 3 × 0.8 ml High Salt Wash Buffer, then eluted into 25 μl streptavidin elution buffer (100 mM DTT, 20 mM HEPES 7.4, 1 mM EDTA, 100 mM NaCl, 0.05% Tween-20) at RT with shaking, then the elution step repeated and combined for a total of 50 μl. At this point, 10% input RNA (4 μl) was diluted into 50 μl streptavidin elution buffer and processed the same as the labeled RNA samples to determine the extent of recovery. 50 μl each input and purified RNA was adjusted to 100 μl with nuclease-free water and purified on RNeasy columns (Qiagen) using the modified protocol as described (Duffy and Simon, 2016). To each 100 μl sample, 350 μl RLT lysis buffer (supplied by the Qiagen kit and supplemented with 10 μl 1% βME per 1 ml RLT) and 250 μl 100% ethanol was added, mixed well, and applied to columns. Columns were washed with 500 μl RPE wash buffer (supplied by the Qiagen kit and supplemented with 35 μl 1% βME per 500 μl RPE), followed by a final 5 min spin at max speed. RNAs were eluted into 14 μl nuclease-free water.

After purification of mRNA, one sample per batch of preps prepared in a single day was tested for enrichment of labeled RNA by RT qPCR, probing both unlabeled and labeled RNA from at least three transcribed genes. The purified 4TU RNA typically contained 2–10% contamination of unlabeled RNA.

### Preparation of 4ThioU RNA libraries for NGS

Newly synthesized RNA isolated via 4-thioU labeling and purification was prepared for sequencing using the Ovation SoLo or Ovation Universal RNA-seq System kits (Tecan) according to the manufacturer’s instructions and 1 ng (SoLo) or 50 ng (Universal) input RNA. Libraries were sequenced on the Illumina HiSeq2500 platform using 25 bp paired-ends at the Fred Hutchinson Genomics Shared Resources facility.

### Immunoprecipitation assays

At OD 0.8-1.0, 1 L cell culture was split into 2 x 500 ml and treated with either DMSO or IAA for 30 min at 30°C. Cells were pelleted and washed in 50 ml Mediator IP buffer (50 mM HEPES, pH 7.6 at 4°C, 150 mM NaCl, 10% glycerol, 0.1% Tween), then washed again in 25 ml buffer containing 1 mM DTT, 1 mM PMSF, 2 ug/ml chymostatin, 0.3 ug/ml leupeptin, 1.4 ug/ml pepstatin, and 0.31 mg/ml benazamidine. Cells were pelleted, frozen in liquid nitrogen, and stored at -80°C until extract preparation. Whole cell extracts were prepared by resuspending cell pellets in ∼1.2 ml buffer + 1 mM DTT + protease inhibitors as described above. Cell suspension was transferred to two 2 ml skirted screw-cap tubes containing 1.25 ml zirconia beads (0.5 mm, BioSpec) and additional buffer added to completely fill tube. Cells were lysed in a Mini Beadbeater-96 (BioSpec Products) five times for 3 min each, with a 1 min chill in ice water in between each lysis step. Cell lysates were spun in an inverted tube for 10 min at 3K rpm after two holes were punctured in the cap using a hot 20 gauge needle. Extracts were transferred to a 1.5 ml microfuge tube and spun at max speed in at 4°C. Supernatant was transferred to new tube, and the final extract was quickly frozen on dry ice and stored at -80°C. Extracts were typically 25-40 mg/ml.

Polyclonal antibodies used in IPs (*α*-Med17, *α*-Med3) were conjugated to Protein G Dynabeads (Life Technologies) in batch as follows, with all incubations and washes performed at room temp. 30 μl Dynabeads per IP reaction were washed twice with PBS, then resuspended in 30 μl PBS and incubated with 10 μl antibody (or 10 μl PBS for beads alone control) with gentle rocking on a Nutator for 30 min. Beads were washed three times with 500 μl PBS, twice with 1 ml 0.2 M triethanolamine pH 8.2, then incubated with 1 ml 0.2 M triethanolamine containing 25 mM dimethyl pimelimidate (DMP) for 30 min on the Nutator. Beads were washed in 500 μl PBS for 5 min. DMP incubation and PBS wash were repeated twice for a total of three times. Beads were washed in 1 ml 0.1 M ethanolamine pH 8.2 for 5 min, followed by a 0.5 ml PBS wash. Beads were washed twice in 1 M glycine pH 3.0 for 10 min each. Finally, beads were washed three times in 1 ml Mediator IP buffer containing 1 mM DTT, protease inhibitors, and 5 mg/ml BSA for 10 min each wash. Beads were resuspended in 30 ul IP buffer containing 1 mM DTT and protease inhibitors.

IPs were performed as follows, with all incubations and washes performed at 4°C. 150-200 ul WCE was incubated with 30 μl antibody-conjugated beads in a final volume of 400 μl IP buffer containing 1 mM DTT and protease inhibitors overnight with gentle rocking on the Nutator. Beads were washed three times with 1 ml Mediator wash buffer (same as IP buffer, except containing 250 mM NaCl) for 10 min each wash. Proteins were eluted in 15 μl 0.1 M glycine pH 2.5 for 10 min at room temp with gentle shaking. Elution was repeated for a total volume of 30 μl. 10 μl 1 M Tris pH 8.5, 3 μl 1 M DTT, and 15 μl 4X LDS sample buffer (Invitrogen) was added to the eluate for a final volume of ∼60 μl. Samples were heated at 70°C for 10 min before running on SDS-PAGE.

### ChEC-seq experiments

ChEC-seq was performed as previously described (Donczew and Hahn, 2021; Donczew et al., 2020b). *S. cerevisiae* 50 ml cultures were pelleted at 2000 x g for 3 min. Cells were resuspended in 1 ml of Buffer A (15 mM Tris, 7.5, 80 mM KCl, 0.1 mM EGTA, 0.2 mM spermine (Millipore Sigma #S3256), 0.3 mM spermidine (Millipore Sigma #85558), protease inhibitors (Millipore Sigma #04693159001)), transferred to a 1.5 ml tube and pelleted at 1500 x g for 30 sec. Cells were washed twice with 1 ml of Buffer A and finally resuspended in 570 μl of Buffer A. 30 μl 2% digitonin (Millipore Sigma #300410) was added to a final concentration of 0.1% and cells were permeabilized for 5 min at 30°C with shaking (900 rpm). 0.2 mM CaCl2 was added to the samples followed by incubation for another 5 min at 30°C. 100 μl cell suspension was mixed with 100 μl Stop Solution (400 mM NaCl, 20 mM EDTA, 4 mM EGTA). Stop Solution was supplemented with 5 μg MNase digested *D. melanogaster* chromatin. Samples were incubated with 0.4 mg/ml Proteinase K (Thermo Fisher Scientific #AM2548) for 30 min at 55°C and the DNA was purified by phenol/CHCl_3_/isoamyl alcohol (25:24:1) extraction and ethanol precipitation. Pellets were resuspended in 30 μl 0.3 mg/ml RNase A (Thermo Fisher Scientific #EN0531) (10 mM Tris, 7.5, 1 mM EDTA, 0.3 mg/ml RNase A) and incubated for 15 min at 37°C. 60 μl of Mag-Bind reagent (Omega Biotek #M1378-01) was added and the samples were incubated for 10 min at RT. Supernatants were transferred to a new tube and the volume was adjusted to 200 μl (10 mM Tris, 8.0, 100 mM NaCl). DNA was purified again by phenol/CHCl_3_/isoamyl alcohol (25:24:1) extraction and ethanol precipitation, and resuspended in 25 μl 10 mM Tris, 8.0.

### ChIP-seq experiments

ChIP-seq experiments were performed similarly as described (Donczew et al., 2020a). 100 ml *S. cerevisiae* or *S. pombe* cultures were crosslinked with 1% formaldehyde (Sigma-Aldrich #252549) for 20 min in the above growth conditions, followed by another 5 min treatment with 130 mM glycine. Cells were pelleted at 3000 x g for 5 min, washed with cold TBS buffer, pelleted at 2000 x g for 3 min, flash-frozen in liquid N_2_, and then stored at −80°C for further use. Cell pellets were resuspended in 300 μl Breaking Buffer (100 mM Tris, 8.0, 20% glycerol, protease inhibitors (Millipore Sigma #04693159001)). Cells were lysed using 0.4 ml zirconia/silica beads (RPI #9834) in a Mini Beadbeater-96 (BioSpec Products) for 5 min. Lysates were spun at 21K x g for 2 min. Pellets were resuspended in 1 ml FA buffer (50 mM HEPES, 7.5, 150 mM NaCl, 1 mM EDTA, 1% Triton X-100, 0.1% sodium deoxycholate, protease inhibitors (Millipore Sigma #04693159001)) and transferred to 15 ml polystyrene tubes. In experiments with antibodies specific against phosphorylated Rpb1 CTD, Breaking Buffer and FA buffer were supplemented with phosphatase inhibitors (Thermo Fisher Scientific #A32957). Samples were sonicated in a cold Bioruptor sonicator bath (Diagenode #UCD-200) at a maximum output, cycling 30 sec on, 30 sec off, for a total of 45 min. Samples were spun twice in fresh tubes at 21K x g for 15 min. Prepared chromatin was flash-frozen in liquid N_2_, and then stored at −80°C for further use.

20 μl of the chromatin sample was used to estimate DNA concentration. First, 20 μl Stop buffer (20 mM Tris, 8.0, 100 mM NaCl, 20 mM EDTA, 1% SDS) was added to samples followed by incubation at 70°C for 16-20 hrs. Samples were digested with 0.5 mg/ml RNase A (Thermo Fisher Scientific #EN0531) for 30 min at 55°C and 1 mg/ml Proteinase K for 90 min at 55°C. Sample volume was brought to 200 μl and DNA was purified by two phenol/CHCl_3_/isoamyl alcohol (25:24:1) extractions and ethanol precipitation. DNA was resuspended in 20 μl 10 mM Tris, 8.0 and the concentration was measured using Qubit HS DNA assay (Thermo Fisher Scientific #Q32851).

20 μl Protein G Dynabeads (Thermo Fisher Scientific #10003D) was used for a single immunoprecipitation. Beads were first washed three times with 500 μl PBST buffer (PBS buffer supplemented with 0.1% Tween 20) for 3 min with gentle rotation. Beads were resuspended in a final volume of 20 μl containing PBST buffer and the indicated antibody. The bead suspension was incubated for 60 min with shaking (1400 rpm) at RT, washed with 500 μl PBST buffer and 500 μl FA buffer. Beads were resuspended in 25 μl FA buffer. 1.5 μg *S. cerevisiae* chromatin and 30 ng *S. pombe* chromatin (strain Sphc821) were combined and samples were brought to a final volume of 500 μl. 25 μl of each sample was mixed with 25 μl Stop buffer and set aside (input sample). 25 μl of beads was added to remaining 475 μl of samples followed by incubation for 16-20 hrs at 4°C.

The beads were washed for 3 min with gentle rotation with the following: 3 times with 500 μl FA buffer, 2 times with FA-HS buffer (50 mM HEPES, 7.5, 500 mM NaCl, 1 mM EDTA, 1% Triton X-100, 0.1% sodium deoxycholate), once with 500 μl RIPA buffer (10 mM Tris, 8.0, 0.25 M LiCl, 0.5% NP-40, 1 mM EDTA, 0.5% sodium deoxycholate). DNA was eluted from beads with 25 μl Stop buffer at 75°C for 10 min. Elution was repeated, eluates were combined and incubated at 70°C for 16-20 hrs together with input samples collected earlier. Samples were digested with 0.5 mg/ml RNase A (Thermo Fisher Scientific #EN0531) for 30 min at 55°C and 1 mg/ml Proteinase K for 2 hrs at 55°C. Sample volume was brought to 200 μl and DNA was purified by two phenol/CHCl_3_/isoamyl alcohol (25:24:1) extractions and ethanol precipitation. DNA was resuspended in 15 μl 10 mM Tris, 8.0 and the concentration was measured using Qubit HS DNA assay (Thermo Fisher Scientific #Q32851).

### Preparation of NGS libraries for ChEC-seq and ChIP-seq samples

NGS libraries for ChEC-seq and ChIP-seq experiments were prepared similarly as described (Donczew et al., 2020a; Warfield et al., 2017). 12 μl of ChEC samples and 5 μl of ChIP samples was used as input for library preparation. Samples were end-repaired, phosphorylated and adenylated in 50 μl reaction volume using the following final concentrations: 1X T4 DNA ligase buffer (NEB #B0202S), 0.5 mM each dNTP (Roche #KK1017), 0.25 mM ATP (NEB #P0756S), 2.5% PEG 4000, 2.5 U T4 PNK (NEB #M0201S), 0.05 U T4 DNA polymerase (Invitrogen #18005025), and 0.05 U Taq DNA polymerase (Thermo Fisher Scientific #EP0401). Reactions were incubated at 12°C 15 min, 37°C 15 min, 72°C 20 min, then put on ice and immediately used in adaptor ligation reactions. Adaptor ligation was performed in a 115 μl volume containing 6.5 nM adaptor, 1X Rapid DNA ligase buffer (Enzymatics #B101L) and 3000 U DNA ligase (Enzymatics #L6030-HC-L) and reactions were incubated at 20 deg for 15 min. Following ligation, a two-step cleanup was performed for ChEC-seq samples using 0.25x vol Mag-Bind reagent (Omega Biotek # M1378-01) in the first step and 1.1x vol in the second step. In case of ChIP-seq samples a single cleanup was performed using 0.4x vol Mag-Bind reagent. In both cases DNA was eluted with 20 μl 10 mM Tris, 8.0. Library Enrichment was performed in a 30 μl reaction volume containing 20 μl DNA from the previous step and the following final concentrations: 1X KAPA buffer (Roche #KK2502), 0.3 mM each dNTP (Roche #KK1017), 2.0 μM each P5 and P7 PCR primer, and 1 U KAPA HS HIFI polymerase (#KK2502). DNA was amplified with the following program: 98°C 45 s, [98°C 15 s, ramp to 60°C @ 3°C /s, 60°C 10 s, ramp to 98°C @ 3°C /s] 16-18x, 72°C 1 min. 18 cycles were used for library amplification for ChEC-seq samples and 16 cycles for ChIP-samples. A post-PCR cleanup was performed using 1.4x vol Mag-Bind reagent and DNA was eluted into 30 μl 10 mM Tris, 8.0. Libraries were sequenced on the Illumina HiSeq2500 platform using 25 bp paired-end reads at the Fred Hutchinson Cancer Research Center Genomics Shared Resources facility.

### Analysis of NGS data

Data analysis was performed similarly as described (Donczew et al., 2020a). The majority of the data analysis tasks except sequence alignment, read counting and peak calling (described below) were performed through interactive work in the Jupyter Notebook (https://jupyter.org) using Python programming language (https://www.python.org) and short Bash scripts. All figures were generated using Matplotlib and Seaborn libraries for Python; (https://matplotlib.org; https://seaborn.pydata.org). All code snippets and whole notebooks are available upon request.

Paired-end sequencing reads were aligned to *S. cerevisiae* reference genome (sacCer3), *S. pombe* reference genome (ASM294v2.20) or *D. melanogaster* reference genome (release 6.06) with Bowtie (Langmead and Salzberg, 2012) using optional arguments ‘-I 10 -X 700 --local -- very-sensitive-local --no-unal --no-mixed --no-discordant’. Details of the analysis pipeline depending on the experimental technique used are described below.

### Analysis of RNA-seq data

SAM files for *S. cerevisiae* data were used as an input for HTseq-count (Anders et al., 2015) with default settings. The GFF file with *S. cerevisiae* genomic features was downloaded from the Ensembl website (assembly R64-1-1). Signal per gene was normalized by the number of all *S. pombe* reads mapped for the sample and multiplied by 10000 (arbitrarily chosen number). Genes classified as dubious, pseudogenes or transposable elements were excluded leaving 5797 genes for the downstream analysis. As a next filtering step, we excluded all the genes that had no measurable signal in at least one out of 168 samples collected in this work under normal growth conditions. The remaining 5283 genes were used to calculate coefficient of variation (CV) to validate reproducibility between replicate experiments (**Figure S1C, S2C and Table S1**). This gene set was further compared to a list of 4900 genes we found previously to provide the best reproducibility with a minimal loss of information (Donczew et al., 2020a). The overlapping set of 4885 genes was used in all plots where genes were divided into Tail-dependent and Tail-independent categories. The results of biological replicate experiments for each sample were averaged. Corresponding samples were compared to calculate log_2_ change in expression per gene (IAA to DMSO samples for degron experiments, Med6_delH1 deletion mutant to Med6_WT strain, Med7_EWE3 mutant to Med7_WT strain, Med21_EWE4 mutant to Med21_WT strain, and ***MED7N, MED15*** and ***MED16*** deletion mutants to med15_med16 _WT strain) (**Table S1**).

### Analysis of ChEC-seq data

SAM files for *S. cerevisiae* data were converted to tag directories with the HOMER (http://homer.ucsd.edu (Heinz et al., 2010)) ‘makeTagDirectory’ tool. Data for three DMSO treated replicate samples from all experiments were used to define bound promoters as described below (**Table S3**). Peaks were called using HOMER ‘findPeaks’ tool with optional arguments set to ‘-o auto -C 0 L 2 F 2’, with the free MNase dataset used as a control. These settings use a default false discovery rate (0.1%) and require peaks to be enriched 2-fold over the control and 2-fold over the local background. Resulting peak files were converted to BED files using ‘pos2bed.pl’ program. For each peak, the peak summit was calculated as a mid-range between peak borders. For peak assignment to promoters the list of all annotated ORF sequences (excluding sequences classified as ‘dubious’ or ‘pseudogene’) was downloaded from the SGD website (https://www.yeastgenome.org). Data for 5888 genes were merged with TSS positions (Park et al., 2014). If the TSS annotation was missing (682 genes), TSS was manually assigned at position −100 bp relative to the start codon. Peaks were assigned to promoters if their peak summit was in the range from −200 to +200 bp relative to TSS for Med10 and Med19 MNase strains or -500 to TSS in strains with MNase fusion on other Mediator subunits. In a few cases, where more than one peak was assigned to the promoter, the one closer to TSS was used. Promoters bound in at least two out of three replicate experiments were included in a final list of promoters bound by a given factor and were used to calculate promoter occupancy in all relevant experiments.

Coverage at each base pair of the *S. cerevisiae* genome was calculated as the number of reads that mapped at that position divided by the number of all *D. melanogaster* reads mapped for the sample and multiplied by 10000 (arbitrarily chosen number). To quantify log_2_ change in factor promoter occupancy signal per promoter was calculated as a sum of normalized reads per base in a 200 bp window around the promoter peak summit. Peak summits were defined using Homer as described above. If no peak was assigned to a promoter using Homer for a given replicate experiment, the position of the strongest signal around TSS was used as a peak summit. Manual inspection of selected cases confirmed the validity of this approach. Corresponding IAA and DMSO treated samples were compared to calculate log_2_ change in occupancy and the results of biological replicate experiments for each sample were averaged (**Table S3**).

### Analysis of ChIP-seq data

For all samples coverage at each base pair of the *S. cerevisiae* genome was calculated as the number of reads that mapped at that position divided by the number of all *S. pombe* reads mapped in the sample, multiplied by the ratio of *S. pombe* to *S. cerevisiae* reads in the corresponding input sample and multiplied by 10000 (arbitrarily chosen number). The list of all annotated ORF sequences (excluding sequences classified as ‘dubious’ or ‘pseudogene’) was downloaded from the SGD website (https://www.yeastgenome.org). Data for 5888 genes were merged with TSS positions obtained from (Park et al., 2014). If the TSS annotation was missing the TSS was manually assigned at position −100 bp relative to the start codon. TFIIB signal per gene was calculated as a sum of normalized reads per base in a fixed window of -200 to +100 relative to TSS (defined in figure legends). The results of biological replicates for each sample were averaged and the log_2_ change in signal per gene was calculated by comparing corresponding IAA and DMSO treated samples (**Table S3**).

## Author contributions

Conceptualization, L.W, R.D., L.M., S.H.; Investigation, L.W., L.M., R.D., S.H.; Formal Analysis, R.D., L.M.; Writing – L.W., R.D., L.M., S.H.; Funding Acquisition and Supervision, S.H.

## Declaration of interests

The authors declare no competing interests.

## Acknowledgements

We thank Olivia Sommers for strain construction and members of the Hahn lab for comments and suggestions throughout the course of this work and J. Schofield for comments on the manuscript. This work was supported by NIH (RO1 GM075114 and R35 GM140823 to SH and P30 CA015704 to the Fred Hutch Genomics and Computational Shared Resources facility.

## Figure and Table Legends

**Table S1. Summary of 4-ThioU RNA-seq experiments**.

**Table S2. List of Med15, Med8, Eaf5, Snf5 and Swr1 binding sites, NDR width and UAS locations of TI and TD genes**

**Table S3. Summary of ChIP-seq and ChEC-seq experiments**.

**Table S4. Yeast strains used in this work**.

**Fig S1. Growth of degron strains, protein depletion and variation in RNA-seq experiments. (A)** Cell growth assays of strains used for RNA-seq. Strains were grown in YPD + adenine overnight at 30°C, diluted to 5 x 10^7^ cells/ml, then 3.5 μl of 10-fold dilutions were spotted to YPD + adenine plates and grown for 2 d at 30°C. **(B)** Western blot analysis of strains treated with DMSO or IAA. Samples for protein analysis taken just prior to 4TU-labeling. Proteins were **s**eparated by SDS-PAGE (4-12% Bis-Tris or 3-8% Tris-Acetate), transferred to PVDF membrane, and visualized by probing with *α*-V5 (degron-tagged subunit) or *α*-Tfg2 (loading control). A titration of DMSO-treated cell extracts was used to determine the extent of protein degradation. **(C)** Boxplot showing coefficients of variation (CV) for all 5283 genes with detectable signals in 168 4-ThioU RNA-seq experiments collected under standard growth conditions. All experiments were done in triplicates except the following samples which were done in duplicates: Med7_EWE3, med15_16_WT, MED15 deletion, MED16 deletion, Med21_EWE4, and Med21_WT. CV values were calculated for each gene based on normalized read counts for replicate experiments.

**Fig S2. Summary of Tail-depletion experiments, transcriptional changes following exposure to stress conditions and properties of CR gene classes. (A)** Scatter plot showing cumulative distribution of log2 change in transcription following TID depletion. Horizontal dashed line indicates the fraction of genes in which transcription changed 1.5-fold. **(B)** Heatmap representation of log2 change in transcription levels in the indicated experiments. Genes are grouped by results of the k-means clustering analysis shown in **Fig. 1A. (C-D)** Boxplots showing dependence on TFIID (Taf1-degron) and Bdf1/2 (Bdf1-degron + *bdf2Δ*;) at the two classes of CR genes: TD (Tail dependent) and TI (Tail-independent). **(E)** Number of promoters bound by Bdf1 (Bdf1 ChEC-seq) at the two classes of CR genes.

**Fig S3. Genome-wide transcription changes in response to stress. (A)** Boxplot showing coefficients of variation (CV) for samples used in stress response analysis. (**B-D**). Scatterplots showing log2 changes in transcription caused by the indicated stress treatment vs expression level under non stress conditions. These results relate to the results in Fig 3 but were calculated based on DMSO treated samples (i.e., Med15 is not depleted).

**Fig S4. Properties of Tail-dependent and Tail-independent genes. (A-E)** Factor binding measured using ChEC-seq (see Methods for details). Used for this analysis are promoters (from the 4885 genes analyzed by RNA-seq in this work) that are bound by a given factor in at least two out of three replicate ChEC or ChIP experiments. (**A**) Bar plot showing percentage of promoters in each class: TD (Tail-dependent) or TI (Tail-independent) bound by MED subunits, Med8 and Med15. (**B**) Bar plot showing the Med8-MNase ChEC-seq signals at UASs of TD and TI genes. (**C-E**) Bar plots showing percentage of promoters in each class bound by: (**C**) SWI/SNF subunit, Snf5. (**D**) NuA4 subunit, Eaf5. (**E**) SWR1 complex subunit, Swr1. (**F**) Box plot showing H4K12-Ac ChIP-seq signal (Donczew and Hahn, 2021) at TD and TI genes. (**G**) Box plot showing the width of nucleosome depleted regions (NDR) (Chereji et al., 2018) at each class. (**H**) Box plot showing the distance between UAS (Median of TFs binding position taken from ChIP-exo data) (Rossi et al., 2021) and TSS (Park et al., 2014) at each class. Welch’s t-test results are shown in **F-H**.

**Fig S5. Mediator structure and the effects of MED mutations on expression of Tail-dependent and independent genes. (A)** Model of yeast MED. Shown is the structure of human MED from the MED-TFIID PIC complex (PDB 7ENC) (Chen et al., 2021) with relevant subunits labeled and showing the Head, Middle, and Tail modules as well as the Plank and Hook domains. Since the complete structure of *S. cerevisiae* MED including the Tail is not yet known, the model approximates yeast MED by omitting the higher eukaryote-specific subunits Med23, 25, 26, 28, 30 leaving subunits conserved between yeasts and humans. **(B)** Boxplot showing log2 change in transcription for 4885 genes measured by 4-thioU RNA-seq after either depleting Med7 N or C-terminal regions or in strains containing the indicated EWE or Med6 mutations. Genes are grouped into Tail-dependent and Tail-independent categories.

**Fig S6. Western blot data for IP results shown in Fig 5.** Extracts were made from cells containing the indicated degrons shown in red after treatment with IAA (+) or DMSO (-). Strains also contained C-terminal 3x Flag epitopes on various MED subunits as indicated. Polyclonal antisera against Med17 (Head), Med3 (Tail) or no antisera (beads alone) were used for immune precipitation. IP eluates were **s**eparated by SDS-PAGE (4-12% Bis-Tris or 3-8% Tris-Acetate), transferred to PVDF membrane, and visualized by probing with rabbit polyclonal antibodies (*α*-Med17, 3) or mouse monoclonal antibodies (*α*-Flag, *α*-V5).

**Fig S7. Representative MED ChEC-seq data and correlations of Pol II, TFIIB and MED binding with transcription.** (A) Genome browser (IGB) images showing comparison of TFIIB ChIP-seq and Med8, Med10, and Med19 ChEC-seq signals at several representative genomic loci. Also shown are Med10-MNase cleavage signals 30 min after degron-depletion of Med14. **(B-C)** Scatterplots showing the correspondence between transcription level and Rpb1 or TFIIB ChIP-seq signals. Results are plotted on log10 scale. Pearson r is shown. Previously published data for RPB1 occupancy were used (Donczew and Hahn, 2021). RPB1 signals were calculated between the annotated TSS and TTS locations (Park et al., 2014). **(D-G)**. Scatterplots showing correlation between MED ChEC-seq signals at core promoters and UAS vs gene expression levels measured by 4ThioU RNA-seq. The gene sets used for this analysis are defined in the text.

## Notes

### Competing Interest Statement

The authors have declared no competing interest.

