## Supplementary Information for "Mediator is broadly recruited to gene promoters via a Tail-independent mechanism"

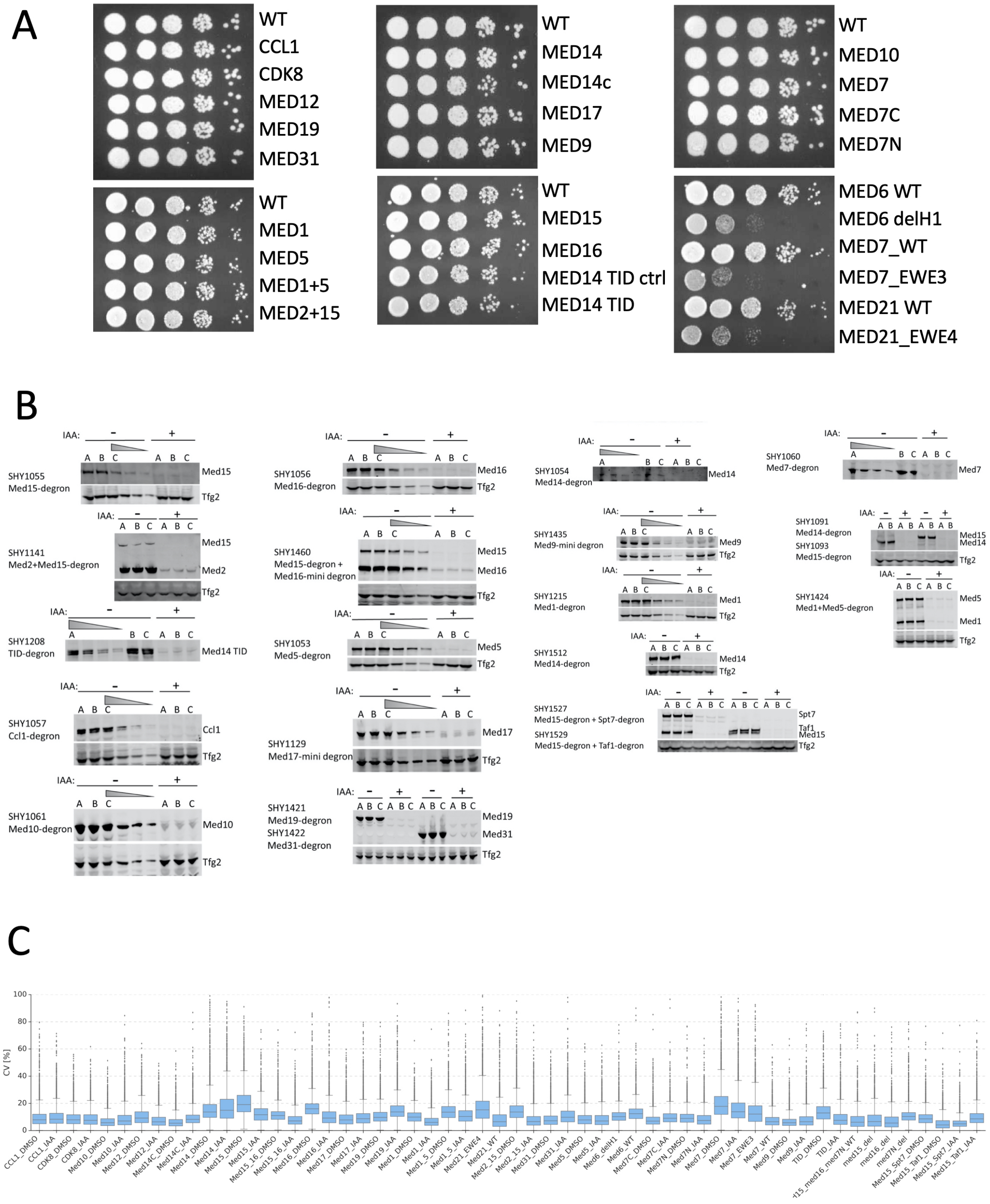


**Fig S1. Growth of degron strains, protein depletion and variation in RNA-seq experiments. (A)** Cell growth assays of strains used for RNA-seq. Strains were grown in YPD + adenine overnight at 30°C, diluted to 5 x $10^7$ cells/ml, then 3.5 µl of 10-fold dilutions were spotted to YPD + adenine plates and grown for 2 d at 30°C. **(B)** Western blot analysis of strains treated with DMSO or IAA. Samples for protein analysis taken just prior to 4TU-labeling. Proteins were **s**eparated by SDS-PAGE (4-12% Bis-Tris or 3-8% Tris-Acetate), transferred to PVDF membrane, and visualized by probing with $\alpha$-V5 (degron-tagged subunit) or $\alpha$-Tfg2 (loading control). A titration of DMSO-treated cell extracts was used to determine the extent of protein degradation. **(C)** Boxplot showing coefficients of variation (CV) for all 5283 genes with detectable signals in 168 4-ThioU RNA-seq experiments collected under standard growth conditions. All experiments were done in triplicates except the following samples which were done in duplicates: Med7_EWE3, med15_16_WT, MED15 deletion, MED16 deletion, Med21_EWE4, and Med21_WT. CV values were calculated for each gene based on normalized read counts for replicate experiments.

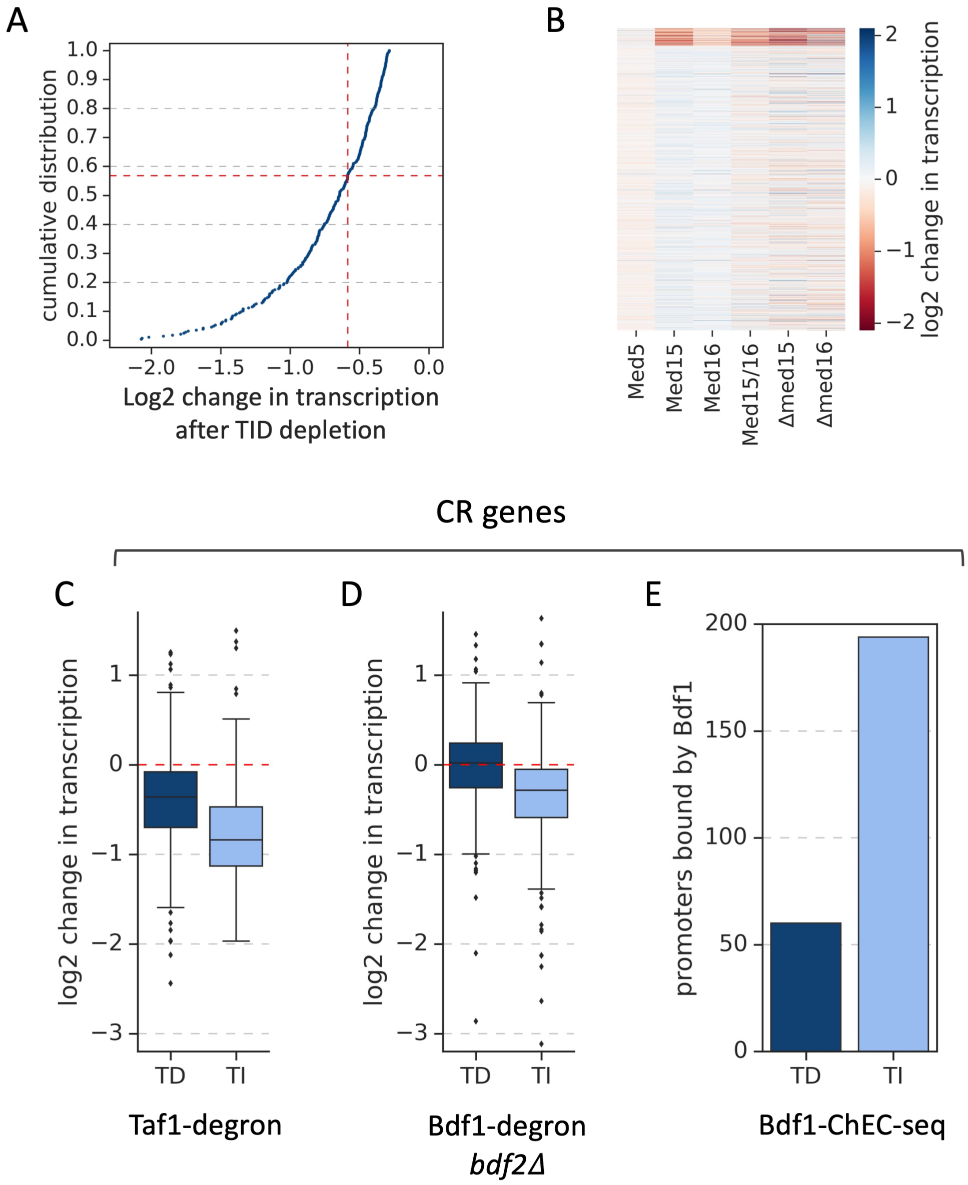


**Fig S2. Summary of Tail-depletion experiments, transcriptional changes following exposure to stress conditions and properties of CR gene classes. (A)** Scatter plot showing cumulative distribution of log2 change in transcription following TID depletion. Horizontal dashed line indicates the fraction of genes in which transcription changed 1.5-fold. **(B)** Heatmap representation of log2 change in transcription levels in the indicated experiments. Genes are grouped by results of the k-means clustering analysis shown in **Fig. 1A. (C-D)** Boxplots showing dependence on TFIID (Taf1-degron) and Bdf1/2 (Bdf1-degron + *bdf2Δ*) at the two classes of CR genes: TD (Tail dependent) and TI (Tail-independent). **(E)** Number of promoters bound by Bdf1 (Bdf1 ChEC-seq) at the two classes of CR genes.

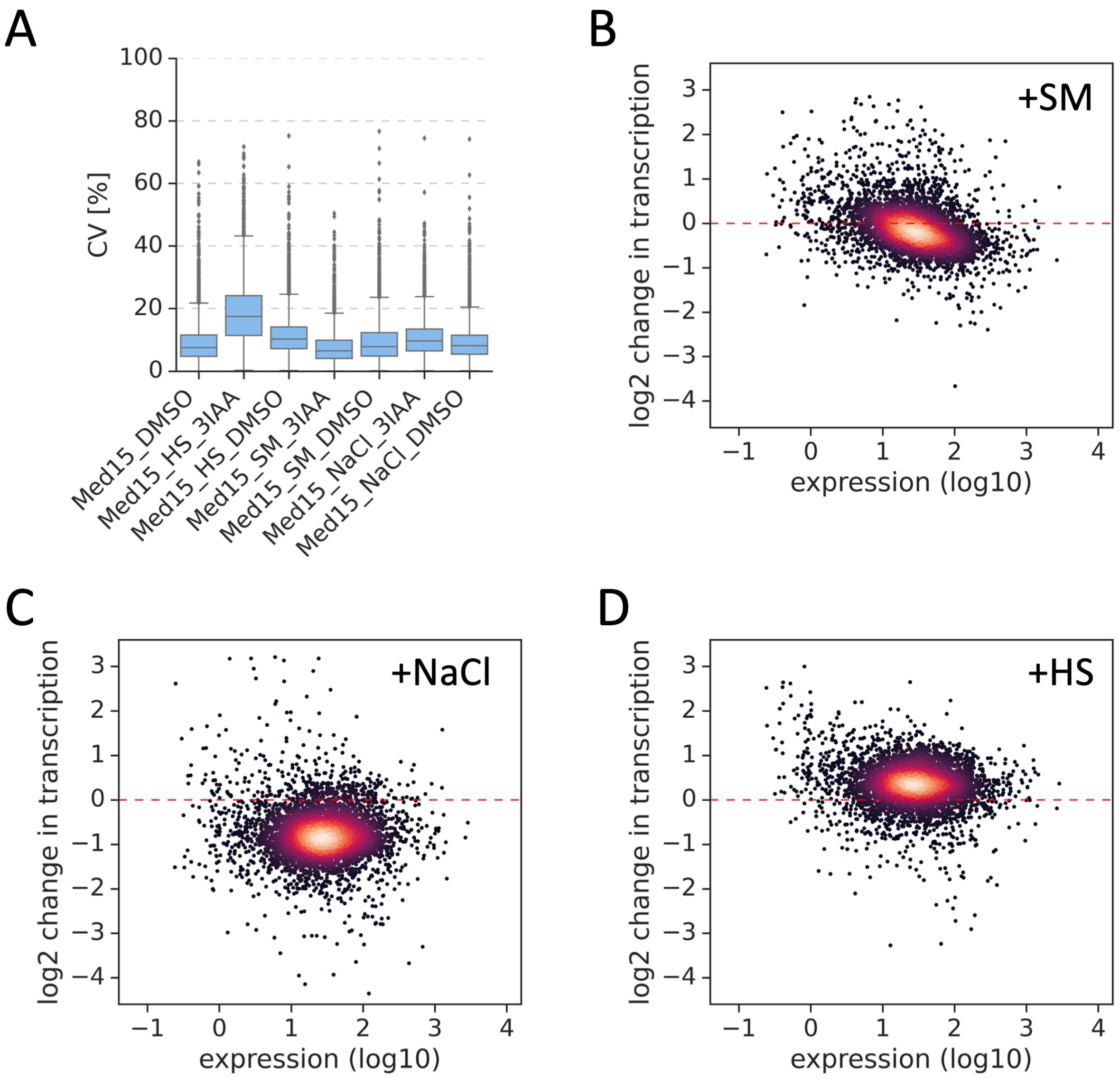


**Fig S3. Genome-wide transcription changes in response to stress. (A)** Boxplot showing coefficients of variation (CV) for samples used in stress response analysis. (**B-D**). Scatterplots showing log2 changes in transcription caused by the indicated stress treatment vs expression level under non stress conditions. These results relate to the results in **Fig 3** but were calculated based on DMSO treated samples (i.e., Med15 is not depleted).

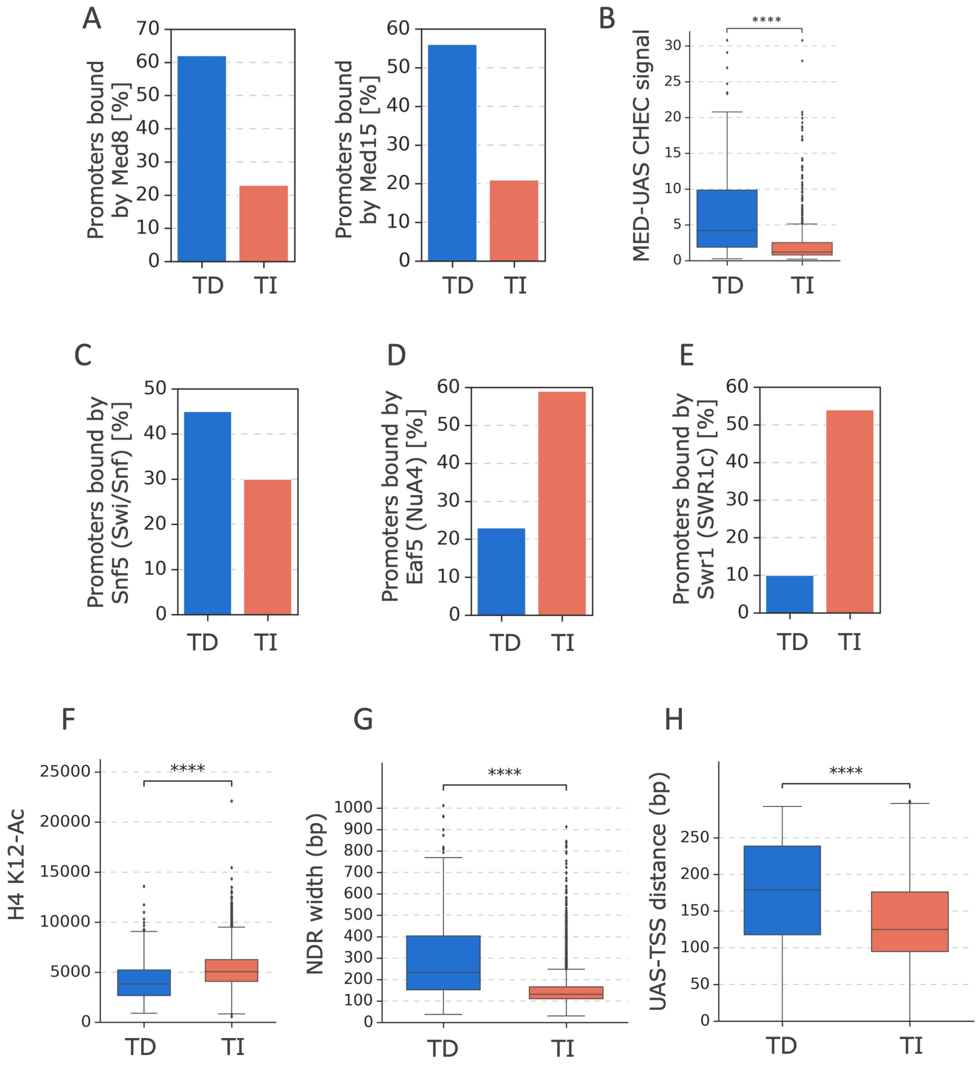


**Fig S4. Properties of Tail-dependent and Tail-independent genes**. (**A-E**) Factor binding measured using ChEC-seq (see Methods for details). Used for this analysis are promoters (from the 4885 genes analyzed by RNA-seq in this work) that are bound by a given factor in at least two out of three replicate ChEC or ChIP experiments. (**A**) Bar plot showing percentage of promoters in each class: TD (Tail-dependent) or TI (Tail-independent) bound by MED subunits, Med8 and Med15. (**B**) Bar plot showing the Med8-MNase ChEC-seq signals at UASs of TD and TI genes. (**C-E**) Bar plots showing percentage of promoters in each class bound by: (**C**) SWI/SNF subunit, Snf5. (**D**) NuA4 subunit, Eaf5. (**E**) SWR1 complex subunit, Swr1. (**F**) Box plot showing H4K12-Ac ChIP-seq signal (Donczew and Hahn, 2021) at TD and TI genes. (**G**) Box plot showing the width of nucleosome depleted regions (NDR) (Chereji et al., 2018) at each class. (**H**) Box plot showing the distance between UAS (Median of TFs binding position taken from ChIP-exo data) (Rossi et al., 2021) and TSS (Park et al., 2014) at each class. Welch's t-test results are shown in **F-H**.

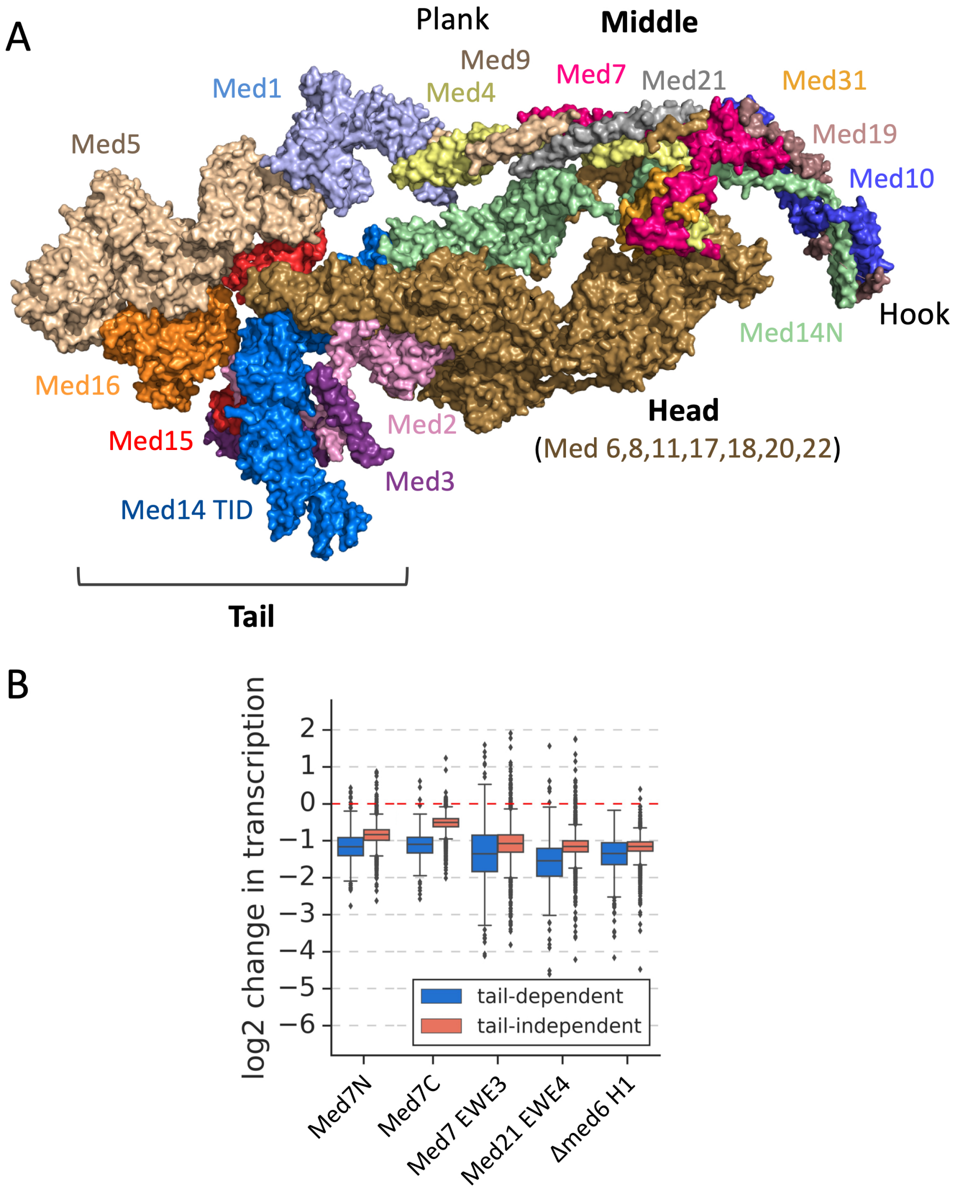


**Fig S5. Mediator structure and the effects of MED mutations on expression of Tail-dependent and independent genes. (A)** Model of yeast MED. Shown is the structure of human MED from the MED-TFIID PIC complex (PDB 7ENC) (Chen et al., 2021) with relevant subunits labeled and showing the Head, Middle, and Tail modules as well as the Plank and Hook domains. Since the complete structure of *S. cerevisiae* MED including the Tail is not yet known, the model approximates yeast MED by omitting the higher eukaryote-specific subunits Med23, 25, 26, 28, 30 leaving subunits conserved between yeasts and humans. **(B)** Boxplot showing log2 change in transcription for 4885 genes measured by 4-thioU RNA-seq after either depleting Med7 N or C-terminal regions or in strains containing the indicated EWE or Med6 mutations. Genes are grouped into Tail-dependent and Tail-independent categories.

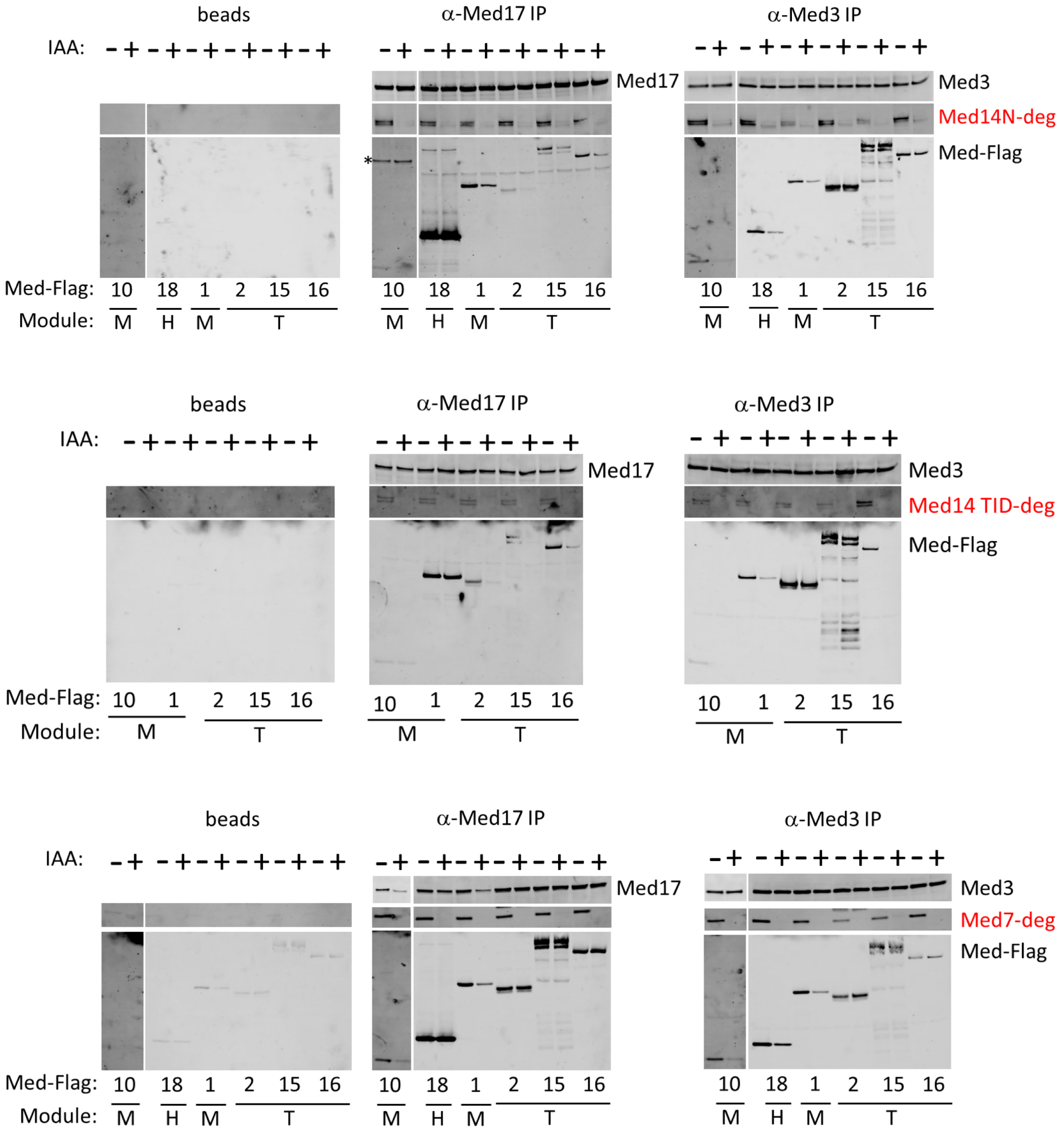


**Fig S6. Western blot data for IP results shown in Fig 5**. Extracts were made from cells containing the indicated degrons shown in red after treatment with IAA (+) or DMSO (-). Strains also contained C-terminal 3x Flag epitopes on various MED subunits as indicated. Polyclonal antisera against Med17 (Head), Med3 (Tail) or no antisera (beads alone) were used for immune precipitation. IP eluates were separated by SDS-PAGE (4-12% Bis-Tris or 3-8% Tris-Acetate), transferred to PVDF membrane, and visualized by probing with rabbit polyclonal antibodies (α-Med17, 3) or mouse monoclonal antibodies (α-Flag, α-V5).

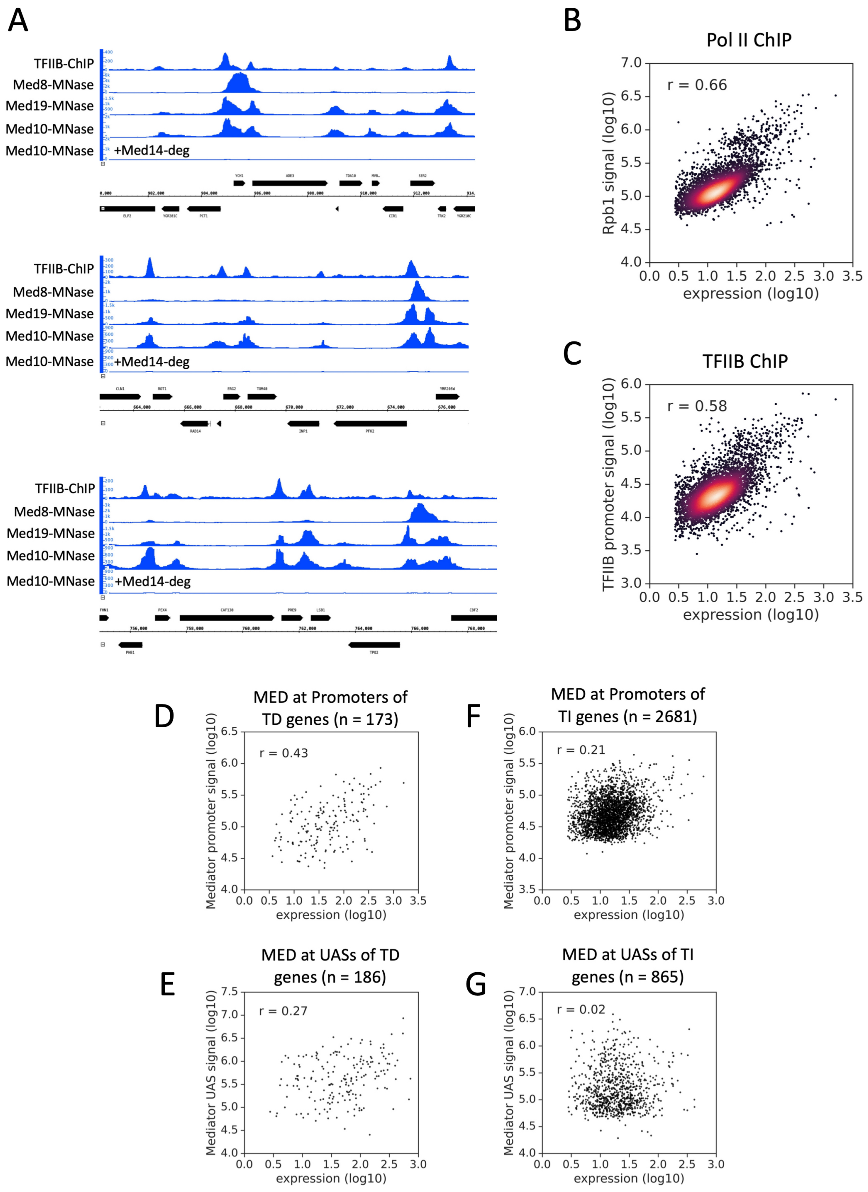


**Fig S7. Representative MED ChEC-seq data and correlations of Pol II, TFIIB and MED binding with transcription. (A)** Genome browser (IGB) images showing comparison of TFIIB ChIP-seq and Med8, Med10, and Med19 ChEC-seq signals at several representative genomic loci. Also shown are Med10-MNase cleavage signals 30 min after degron-depletion of Med14. **(B-C)** Scatterplots showing the correspondence between transcription level and Rpb1 or TFIIB ChIP-seq signals. Results are plotted on log10 scale. Pearson r is shown. Previously published data for RPB1 occupancy were used (Donczew and Hahn, 2021). RPB1 signals were calculated between the annotated TSS and TTS locations (Park et al., 2014). **(D-G)**. Scatterplots showing correlation between MED ChEC-seq signals at core promoters and UAS vs gene expression levels measured by 4ThioU RNA-seq. The gene sets used for this analysis are defined in the text.

**Supplementary Figures References**

Chen, X., Yin, X., Li, J., Wu, Z., Qi, Y., Wang, X., Liu, W., and Xu, Y. (2021). Structures of the human Mediator and Mediator-bound preinitiation complex. Science *372*, eabg0635.

Chereji, R.V., Ramachandran, S., Bryson, T.D., and Henikoff, S. (2018). Precise genome-wide mapping of single nucleosomes and linkers in vivo. Genome Biol *19*, 19.

Donczew, R., and Hahn, S. (2021). BET family members Bdf1/2 modulate global transcription initiation and elongation in Saccharomyces cerevisiae. Elife *10*, e69619.

Park, D., Morris, A.R., Battenhouse, A., and Iyer, V.R. (2014). Simultaneous mapping of transcript ends at single-nucleotide resolution and identification of widespread promoter-associated non-coding RNA governed by TATA elements. Nucleic Acids Research *42*, 3736–3749.

Rossi, M.J., Kuntala, P.K., Lai, W.K.M., Yamada, N., Badjatia, N., Mittal, C., Kuzu, G., Bocklund, K., Farrell, N.P., Blanda, T.R., et al. (2021). A high-resolution protein architecture of the budding yeast genome. Nature *592*, 309–314.

**Supplementary Table 4:** Yeast Strains

| Strain | Description | Genotype | Reference |
|---|---|---|---|
| BY4705 | Wild type (WT) | *MATα Δade2::hisG his3Δ200 leu2Δ0 lys2Δ0 met15Δ0 trp1Δ63 ura3Δ0* | (Brachmann et al., 1998) |
| SHY1055 | *RPB3*-Flag, *MED15*-degron | *MATα Δade2::hisG his3Δ200 leu2Δ0 lys2Δ0 met15Δ0 trp1Δ63 ura3Δ0 RPB3*-3xFlag::*NATMX pGPD1-OSTIR::HIS3 MED15*-3xV5 IAA7::*KanMX* | This work |
| SHY1141 | *RPB3*-Flag, *MED2+MED15*-degron | *MATα Δade2::hisG his3Δ200 leu2Δ0 lys2Δ0 met15Δ0 trp1Δ63 ura3Δ0 RPB3*-3xFlag::*NATMX pGPD1-OSTIR::HIS3 MED15*-3xV5 IAA7::*KanMX MED2*-3xV5-IAA7::*URA3* | This work |
| SHY1208 | *MED14* TID-degron | *MATα Δade2::hisG his3Δ200 leu2Δ0 lys2Δ0 met15Δ0 trp1Δ63 ura3Δ0 med14ΔORF::KanMX pGPD1-OSTIR::HIS3* pSH1933 (ars cen *TRP1 pRGR1-med14*ΔTID) pSH1940 (ars cen *LEU2 pRGR1-med14* TID-3xV5-degron) | This work |
| SHY1056 | *RPB3*-Flag, *MED16*-degron | *MATα Δade2::hisG his3Δ200 leu2Δ0 lys2Δ0 met15Δ0 trp1Δ63 ura3Δ0 RPB3*-3xFlag::*NATMX pGPD1-OSTIR::HIS3 MED16*-3xV5 IAA7::*KanMX* | This work |
| SHY1460 | *MED15*-degron + *MED16*-mini degron | *MATα Δade2::hisG his3Δ200 leu2Δ0 lys2Δ0 met15Δ0 trp1Δ63 ura3Δ0 pGPD1-OSTIR::HIS3 MED15*-3xV5-IAA7::*URA3 MED16*-3xV5 IAA7 mini degron::*KanMX* | This work |
| SHY825 | *med15Δ* | *MATα Δade2::hisG Δ200 leu2Δ0 lys2Δ0 met15Δ0 trp1Δ63 ura3Δ0 Δgal11::HPH* | This work |
| SHY331 | *med16Δ* | *MATα Δade2::hisG his3Δ200 leu2Δ0 lys2Δ0 met15Δ0 trp1Δ63 ura3Δ0 Δsin4::TRP1* | (Reeves and Hahn, 2003) |

| | | | |
|---|---|---|---|
| SHY1053 | *RPB3*-Flag, *MED5*-degron | *MAT*$\alpha$ *Δade2::hisG his3Δ200 leu2Δ0 lys2Δ0 met15Δ0 trp1Δ63 ura3Δ0 RPB3*-3xFlag::*NATMX pGPD1-OSTIR::HIS3 MED5*-3xV5 IAA7::*KanMX* | This work |
| SHY1057 | *RPB3*-Flag, *CCL1*-degron | *MAT*$\alpha$ *Δade2::hisG his3Δ200 leu2Δ0 lys2Δ0 met15Δ0 trp1Δ63 ura3Δ0 RPB3*-3xFlag::*NATMX pGPD1-OSTIR::HIS3 CCL1*-3xV5 IAA7::*KanMX* | This work |
| SHY1061 | *RPB3*-Flag, *MED10*-degron | *MAT*$\alpha$ *Δade2::hisG his3Δ200 leu2Δ0 lys2Δ0 met15Δ0 trp1Δ63 ura3Δ0 RPB3*-3xFlag::*NATMX pGPD1-OSTIR::HIS3 MED10*-3xV5 IAA7::*KanMX* | This work |
| SHY1054 | *RPB3*-Flag, *MED14*-degron | *MAT*$\alpha$ *Δade2::hisG his3Δ200 leu2Δ0 lys2Δ0 met15Δ0 trp1Δ63 ura3Δ0 RPB3*-3xFlag::*NATMX pGPD1-OSTIR::HIS3 MED14*-3xV5 IAA7::*KanMX* | This work |
| SHY1129 | *RPB3*-Flag, *MED17*-mini degron | *MAT*$\alpha$ *Δade2::hisG his3Δ200 leu2Δ0 lys2Δ0 met15Δ0 trp1Δ63 ura3Δ0 RPB3*-3xFlag::*NATMX pGPD1-OSTIR::HIS3 MED17*-3xV5 IAA7 mini degron::*KanMX* | This work |
| SHY1421 | *MED19*-degron | *MAT*$\alpha$ *Δade2::hisG his3Δ200 leu2Δ0 lys2Δ0 met15Δ0 trp1Δ63 ura3Δ0 pGPD1-OSTIR::HIS3 MED19*-3xV5 IAA7::*KanMX* | This work |
| SHY1422 | *MED31*-degron | *MAT*$\alpha$ *Δade2::hisG his3Δ200 leu2Δ0 lys2Δ0 met15Δ0 trp1Δ63 ura3Δ0 pGPD1-OSTIR::HIS3 MED19*-3xV5 IAA7::*KanMX* | This work |
| SHY1060 | *RPB3*-Flag, *MED7*-degron | *MAT*$\alpha$ *Δade2::hisG his3Δ200 leu2Δ0 lys2Δ0 met15Δ0 trp1Δ63 ura3Δ0 RPB3*-3xFlag::*NATMX pGPD1-OSTIR::HIS3 MED7*-3xV5 IAA7::*KanMX* | This work |
| SHY1435 | *MED9*-mini degron | *MAT*$\alpha$ *Δade2::hisG his3Δ200 leu2Δ0 lys2Δ0 met15Δ0 trp1Δ63 ura3Δ0 pGPD1-OSTIR::HIS3* | This work |

| | | | |
|---|---|---|---|
| | | *MED9*-3xV5 IAA7 mini degron::*KanMX* | |
| SHY1215 | *RPB3*-Flag, *MED1*-degron | *MATα Δade2::hisG his3Δ200 leu2Δ0 lys2Δ0 met15Δ0 trp1Δ63 ura3Δ0 RPB3*-3xFlag::*NATMX pGPD1-OSTIR::HIS3* *MED1*-3xV5 IAA7::*KanMX* | This work |
| SHY1424 | *RPB3*-Flag, *MED1+MED5*-degron | *MATα Δade2::hisG his3Δ200 leu2Δ0 lys2Δ0 met15Δ0 trp1Δ63 ura3Δ0 RPB3*-3xFlag::*NATMX pGPD1-OSTIR::HIS3* *MED1*-3xV5 IAA7::*KanMX MED5*-3xV5 IAA7::*URA3* | This work |
| SHY1091 | *RPB3*-Flag, *SUA7*-Myc, *MED14*-degron | *MATα Δade2::hisG his3Δ200 leu2Δ0 lys2Δ0 met15Δ0 trp1Δ63 ura3Δ0 RPB3*-3xFlag::*NATMX pGPD1-OSTIR::HIS3 SUA7-13x Myc::HPH* *MED14*-3xV5 IAA7::*KanMX* | This work |
| SHY1093 | *RPB3*-Flag, *SUA7*-Myc, *MED15*-degron | *MATα Δade2::hisG his3Δ200 leu2Δ0 lys2Δ0 met15Δ0 trp1Δ63 ura3Δ0 RPB3*-3xFlag::*NATMX pGPD1-OSTIR::HIS3 SUA7-13x Myc::HPH* *MED15*-3xV5 IAA7::*KanMX* | This work |
| SHY1389 | *MED1*-Flag, *MED14N*-degron | *MATα Δade2::hisG his3Δ200 leu2Δ0 lys2Δ0 met15Δ0 trp1Δ63 ura3Δ0 med14ΔORF::KanMX pGPD1-OSTIR::HIS3 MED1*-3xFlag::*NATMX* pSH1936 (ars cen *LEU2 pRGR1-MED14* TID) pSH1957 (ars cen *TRP1 pRGR1-MED14(*Δ TID2)-3xV5 mini degron) | This work |
| SHY1390 | *MED2*-Flag, *MED14N*-degron | *MATα Δade2::hisG his3Δ200 leu2Δ0 lys2Δ0 met15Δ0 trp1Δ63 ura3Δ0 rgr1/med14ΔORF::KanMX pGPD1-OSTIR::HIS3 MED2*-3xFlag::*NATMX* pSH1936 (ars cen *LEU2 pRGR1-MED14* TID) pSH1957 (ars cen *TRP1 pRGR1-* | This work |

| | | | |
|---|---|---|---|
| | | *MED14(*Δ TID2)-3xV5 mini degron) | |
| SHY1391 | *MED10*-Flag, *MED14N*-degron | *MATα Δade2::hisG his3Δ200 leu2Δ0 lys2Δ0 met15Δ0 trp1Δ63 ura3Δ0 rgr1/med14ΔORF::KanMX pGPD1-OSTIR::HIS3 MED10*-3xFlag::*NATMX* pSH1936 (ars cen *LEU2 pRGR1-MED14* TID) pSH1957 (ars cen *TRP1 pRGR1-MED14(*Δ TID2)-3xV5 mini degron) | This work |
| SHY1392 | *MED15*-Flag, *MED14N*-degron | *MATα Δade2::hisG his3Δ200 leu2Δ0 lys2Δ0 met15Δ0 trp1Δ63 ura3Δ0 rgr1/med14ΔORF::KanMX pGPD1-OSTIR::HIS3 MED15*-3xFlag::*NATMX* pSH1936 (ars cen *LEU2* pSH1936 (ars cen *LEU2 pRGR1-MED14* TID) pSH1957 (ars cen *TRP1 pRGR1-MED14(*Δ TID2)-3xV5 mini degron) | This work |
| SHY1393 | *MED16*-Flag, *MED14N*-degron | *MATα Δade2::hisG his3Δ200 leu2Δ0 lys2Δ0 met15Δ0 trp1Δ63 ura3Δ0 rgr1/med14ΔORF::KanMX pGPD1-OSTIR::HIS3 MED16*-3xFlag::*NATMX* pSH1936 (ars cen *LEU2 pRGR1-MED14* TID) pSH1957 (ars cen *TRP1 pRGR1-MED14(*Δ TID2)-3xV5 mini degron) | This work |
| SHY1394 | *MED18*-Flag, *MED14N*-degron | *MATα Δade2::hisG his3Δ200 leu2Δ0 lys2Δ0 met15Δ0 trp1Δ63 ura3Δ0 rgr1/med14ΔORF::KanMX pGPD1-OSTIR::HIS3 MED18*-3xFlag::*NATMX* pSH1936 (ars cen *LEU2 pRGR1-MED14* TID) pSH1957 (ars cen *TRP1 pRGR1-MED14(*Δ TID2)-3xV5 mini degron) | This work |

| | | | |
|---|---|---|---|
| SHY1418 | *MED1*-Flag, *MED14 TID*-degron | *MATα Δade2::hisG his3Δ200 leu2Δ0 lys2Δ0 met15Δ0 trp1Δ63 ura3Δ0 rgr1/med14ΔORF::KanMX pGPD1-OSTIR::HIS3 MED1*-3xFlag::*NATMX* pSH1933 (ars cen *TRP1 pRGR1-MED14*ΔTID) pSH1940 (ars cen *LEU2 pRGR1-RGR1/MED14* TID-3xV5 degron) | This work |
| SHY1385 | *MED2*-Flag, *MED14 TID*-degron | *MATα Δade2::hisG his3Δ200 leu2Δ0 lys2Δ0 met15Δ0 trp1Δ63 ura3Δ0 rgr1/med14ΔORF::KanMX pGPD1-OSTIR::HIS3 MED2*-3xFlag::*NATMX* pSH1933 (ars cen *TRP1 pRGR1-MED14*ΔTID) pSH1940 (ars cen *LEU2 pRGR1-RGR1/MED14* TID-3xV5 degron) | This work |
| SHY1386 | *MED10*-Flag, *MED14 TID*-degron | *MATα Δade2::hisG his3Δ200 leu2Δ0 lys2Δ0 met15Δ0 trp1Δ63 ura3Δ0 rgr1/med14ΔORF::KanMX pGPD1-OSTIR::HIS3 MED10*-3xFlag::*NATMX* pSH1933 (ars cen *TRP1 pRGR1-MED14*ΔTID) pSH1940 (ars cen *LEU2 pRGR1-RGR1/MED14* TID-3xV5 degron) | This work |
| SHY1387 | *MED15*-Flag, *MED14 TID*-degron | *MATα Δade2::hisG his3Δ200 leu2Δ0 lys2Δ0 met15Δ0 trp1Δ63 ura3Δ0 rgr1/med14ΔORF::KanMX pGPD1-OSTIR::HIS3 MED15*-3xFlag::*NATMX* pSH1933 (ars cen *TRP1 pRGR1-MED14*ΔTID) pSH1940 (ars cen *LEU2 pRGR1-RGR1/MED14* TID-3xV5 degron) | This work |
| SHY1388 | *MED16*-Flag, *MED14 TID*-degron | *MATα Δade2::hisG his3Δ200 leu2Δ0 lys2Δ0 met15Δ0 trp1Δ63 ura3Δ0 rgr1/med14ΔORF::KanMX pGPD1-OSTIR::HIS3 MED16*-3xFlag::*NATMX* pSH1933 (ars cen | This work |

| | | | |
|---|---|---|---|
| | | *TRP1 pRGR1-MED14*ΔTID) pSH1940 (ars cen *LEU2 pRGR1-RGR1/MED14* TID-3xV5 degron) | |
| SHY1412 | *MED1*-Flag, *MED7*-degron | *MATα Δade2::hisG his3Δ200 leu2Δ0 lys2Δ0 met15Δ0 trp1Δ63 ura3Δ0 pGPD1-OSTIR::HIS3 MED7*-3xV5 IAA7::*KanMX MED1-3xFlag::NATMX* | This work |
| SHY1413 | *MED2*-Flag, *MED7*-degron | *MATα Δade2::hisG his3Δ200 leu2Δ0 lys2Δ0 met15Δ0 trp1Δ63 ura3Δ0 pGPD1-OSTIR::HIS3 MED7*-3xV5 IAA7::*KanMX MED2-3xFlag::NATMX* | This work |
| SHY1414 | *MED10*-Flag, *MED7*-degron | *MATα Δade2::hisG his3Δ200 leu2Δ0 lys2Δ0 met15Δ0 trp1Δ63 ura3Δ0 pGPD1-OSTIR::HIS3 MED7*-3xV5 IAA7::*KanMX MED10-3xFlag::NATMX* | This work |
| SHY1415 | *MED15*-Flag, *MED7*-degron | *MATα Δade2::hisG his3Δ200 leu2Δ0 lys2Δ0 met15Δ0 trp1Δ63 ura3Δ0 pGPD1-OSTIR::HIS3 MED7*-3xV5 IAA7::*KanMX MED15-3xFlag::NATMX* | This work |
| SHY1416 | *MED16*-Flag, *MED7*-degron | *MATα Δade2::hisG his3Δ200 leu2Δ0 lys2Δ0 met15Δ0 trp1Δ63 ura3Δ0 pGPD1-OSTIR::HIS3 MED7*-3xV5 IAA7::*KanMX MED16-3xFlag::NATMX* | This work |
| SHY1417 | *MED18*-Flag, *MED7*-degron | *MATα Δade2::hisG his3Δ200 leu2Δ0 lys2Δ0 met15Δ0 trp1Δ63 ura3Δ0 pGPD1-OSTIR::HIS3 MED7*-3xV5 IAA7::*KanMX MED18-3xFlag::NATMX* | This work |
| SHY1462 | *med16Δ* | *MATα Δade2::hisG his3Δ200 leu2Δ0 lys2Δ0 met15Δ0 trp1Δ63 ura3Δ0 med16Δ::KanMX* | This work |
| SHY1161 | *MED8*-MNase | *MATα Δade2::hisG his3Δ200 leu2Δ0 lys2Δ0 met15Δ0 trp1Δ63 ura3Δ0 MED8*-MNase::*TRP1* | This work |
| SHY1475 | *MED19*-Flag-MNase, *KIN28* is | *MATα Δade2::hisG his3Δ200 leu2Δ0 lys2Δ0 met15Δ0 trp1Δ63* | This work |

| | | | |
|---|---|---|---|
| | | *ura3Δ0 kin28is MED19*-3xFlag-MNase::*TRP1* | |
| SHY1474 | *MED10*-Flag-MNase, *KIN28* is | *MATα Δade2::hisG his3Δ200 leu2Δ0 lys2Δ0 met15Δ0 trp1Δ63 ura3Δ0 kin28is MED10*-3xFlag-MNase::*TRP1* | This work |
| SHY1512 | *MED10*-Flag-MNase, *MED14*-degron | *MATα Δade2::hisG his3Δ200 leu2Δ0 lys2Δ0 met15Δ0 trp1Δ63 ura3Δ0 pGPD1-OSTIR::HIS3 MED14*-3xV5-IAA7::*KanMX MED10*-3xFlag-MNase::*URA3* | This work |
| SHY1436 | *MED7N*-degron | *MATα Δade2::hisG his3Δ200 leu2Δ0 lys2Δ0 met15Δ0 trp1Δ63 ura3Δ0 pGPD1-OSTIR::HIS3 med7Δ*::*KanMX* pSH1991 *(ars cen LEU2 mini degron-med7ΔC*) pSH1986 (*ars cen TRP1 med7ΔN)* | This work |
| SHY1428 | *MED7C*-degron | *MATα Δade2::hisG his3Δ200 leu2Δ0 lys2Δ0 met15Δ0 trp1Δ63 ura3Δ0 pGPD1-OSTIR::HIS3 med7Δ*::*KanMX* pSH1989 *(ars cen LEU2 med7ΔC*) pSH1990 (*ars cen TRP1 med7ΔN*-mini degron*)* | This work |
| SHY1444 | *MED7* WT | *MATα Δade2::hisG his3Δ200 leu2Δ0 lys2Δ0 met15Δ0 trp1Δ63 ura3Δ0 pGPD1-OSTIR::HIS3 med7Δ*::*KanMX* pSH1970 (*ars cen LEU2 MED7*) | This work |
| SHY1445 | *MED7* EWE3 | *MATα Δade2::hisG his3Δ200 leu2Δ0 lys2Δ0 met15Δ0 trp1Δ63 ura3Δ0 pGPD1-OSTIR::HIS3 med7Δ*::*KanMX* pSH1998 (*ars cen LEU2 med7* EWE3) | This work |
| SHY1442 | *MED21* WT | *MATα Δade2::hisG his3Δ200 leu2Δ0 lys2Δ0 met15Δ0 trp1Δ63 ura3Δ0 pGPD1-OSTIR::HIS3 med21Δ*::*KanMX* pSH1997 *(ars cen LEU2 MED21)* | This work |
| SHY1443 | *MED21* EWE4 | *MATα Δade2::hisG his3Δ200 leu2Δ0 lys2Δ0 met15Δ0 trp1Δ63 ura3Δ0 pGPD1-OSTIR::HIS3* | This work |

| | | | |
|---|---|---|---|
| | | *med21Δ*::*KanMX* pSH2001 *(ars cen LEU2 med21* EWE4*)* | |
| SHY1454 | *MED6* WT | *MATα Δade2::hisG his3Δ200 leu2Δ0 lys2Δ0 met15Δ0 trp1Δ63 ura3Δ0 pGPD1-OSTIR::HIS3 med6Δ*::*KanMX* pSH2008 *(ars cen LEU2 MED6)* | This work |
| SHY1455 | *MED6* ΔH1 | *MATα Δade2::hisG his3Δ200 leu2Δ0 lys2Δ0 met15Δ0 trp1Δ63 ura3Δ0 pGPD1-OSTIR::HIS3 med6Δ*::*KanMX* pSH2009 *(ars cen LEU2 med6Δ Helix 1 residues 13-20)* | This work |
| Sphc821 | *S. pombe* WT / *ABP1*-Flag, *CBH1*-Myc | *S. pombe* h+ leu1-32 ade6-216 *OtrR1:ura4 Abp1*-3xFLAG::*KanMx* Cbh1-13xMyc::*NanMX* | (Cam et al., 2008) From T. Tsukiyama - Fred Hutch |
| SHY1058 | *S. pombe* WT / *RPB3*-Flag | *S. pombe* mating type h- *RPB3*-3xFlag::*KanMX* | From G. Smith lab - Fred Hutch |
| SHY1221 | *EAF5*-MNase | *matα Δade2::hisG his3 Δ200 leu2 Δ0 lys2 Δ0 met15 Δ0 trp1 Δ63 ura3 Δ0EAF5-*MNase::*TRP1* | This work |
| SHY1224 | *SNF5*-MNase | *matα Δade2::hisG his3 Δ200 leu2 Δ 0 lys2 Δ0 met15 Δ0 trp1 Δ63 ura3 Δ0 SNF5*-MNase::*TRP1* | This work |
| SHY1229 | *MED15*-MNase | *matα Δade2::hisG his3 Δ200 leu2 Δ0 lys2 Δ0 met15 Δ0 trp1 Δ63 ura3 Δ0 MED15-*MNase::*TRP1* | This work |
| SHY1342 | *SWR1*-MNase | *matα Δade2::hisG his3 Δ200 leu2 Δ 0 lys2 Δ0 met15 Δ0 trp1 Δ63 ura3 Δ 0 SWR1*-MNase::*TRP1* | This work |
| SHY1527 | *MED15*-degron + *SPT7*-degron | *MATα Δade2::hisG his3Δ200 leu2Δ0 lys2Δ0 met15Δ0 trp1Δ63 ura3Δ0 pGPD1-OSTIR::HIS3 MED15*-3xV5 IAA7::*KanMX SPT7-*3xV5-IAA7::*LEU2* | This work |
| SHY1529 | *MED15*-degron + *TAF1*-degron | *MATα Δade2::hisG his3Δ200 leu2Δ0 lys2Δ0 met15Δ0 trp1Δ63* | This work |

| | | | |
|---|---|---|---|
| | | *ura3Δ0 pGPD1-OSTIR::HIS3 MED15*-3xV5 IAA7::*KanMX TAF1*-3xV5-IAA7::*LEU2* | |
| SHY1176 | *RPB3*-Flag, *SPT3 + SPT7*-degron | *matα Δade2::hisG his3Δ200 leu2Δ0 lys2Δ0 met15Δ0 trp1Δ63 ura3Δ0 RPB3*-3x Flag::*NatMX pGPD1-OSTIR::HIS3 SPT7*-3xV5-IAA7::*KanMX SPT3*-3xV5-IAA7:: *URA3* | (Donczew et al., 2020) |
| SHY1039 | *RPB3*-Flag, *TAF1*-degron | *matα Δade2::hisG his3Δ200 leu2Δ0 lys2Δ0 met15Δ0 trp1Δ63 ura3Δ0 RPB3*-3x Flag::*NatMX pGPD1-OSTIR::HIS3 TAF1*-3xV5-IAA7::*KanMX* | (Donczew et al., 2020) |
| SHY1043 | *RPB3*-Flag, *TAF13*-degron | *matα Δade2::hisG his3Δ200 leu2Δ0 lys2Δ0 met15Δ0 trp1Δ63 ura3Δ0 RPB3*-3x Flag::*NatMX pGPD1-OSTIR::HIS3 TAF13*-3xV5-IAA7::*KanMX* | (Donczew et al., 2020) |
| SHY1291 | *RPB3*-Flag, *TAF13 + SPT7*-degron | *matα Δade2::hisG his3Δ200 leu2Δ0 lys2Δ0 met15Δ0 trp1Δ63 ura3Δ0 RPB3*-3x Flag::*NatMX pGPD1-OSTIR::HIS3 TAF13*-3xV5-IAA7::*KanMX SPT7*-3xV5-IAA7-degron::*URA3* | (Donczew et al., 2020) |
| SHY1213 | *SSL2-mini* degron | *matα Δade2::hisG his3Δ200 leu2Δ0 lys2Δ0 met15Δ0 trp1Δ63 ura3Δ0 rad25Δ::KanMX pGPD1-OSTIR::HIS3* pSH1947 (ars cen LEU2 3xV5-degron3-Ssl2/Rad25) | (Donczew and Hahn, 2021) |
| SHY1306 | *RPB3*-Flag, *MED8*-MNase, MED14-degron | *MATα Δade2::hisG his3Δ200 leu2Δ0 lys2Δ0 met15Δ0 trp1Δ63 ura3Δ0 RPB3*-3xFlag::*NatMX pGPD1-OSTIR::HIS3 MED14-3xV5 IAA7::KanMX MED8-MNase::TRP1* | This work |
| SHY1307 | *RPB3*-Flag, *MED15*-MNase, *MED14*-degron | *MATα Δade2::hisG his3Δ200 leu2Δ0 lys2Δ0 met15Δ0 trp1Δ63 ura3Δ0 RPB3*-3xFlag::*NATMX pGPD1-OSTIR::HIS3 MED14-3xV5 IAA7::KanMX MED15-MNase::TRP1* | This work |

| | | | |
|---|---|---|---|
| SHY1512 | *MED10*-Flag-MNase, *MED14-degron* | *MATα Δade2::hisG his3Δ200 leu2Δ0 lys2Δ0 met15Δ0 trp1Δ63 ura3Δ0 pGPD1-OSTIR::HIS3 MED14*-3xV5-IAA7::*KanMX MED10*-3xFlag-MNase::*URA3* | This work |
| SHY1354 | *MED17*-Flag-MNase, *MED14* TID-degron | *MATα Δade2::hisG his3Δ200 leu2Δ0 lys2Δ0 met15Δ0 trp1Δ63 ura3Δ0 med14ΔORF::KanMX pGPD1-OSTIR::HIS3 MED17-3xFlag-MNase::URA3* pSH1933 (ars cen *TRP1 pRGR1-med14Δ*TID) pSH1940 (ars cen *LEU2 pRGR1-med14 TID-3xV5-degron)* | This work |
| SHY1503 | *MED10*-Flag-MNase, *MED14* TID-degron | *MATα Δade2::hisG his3Δ200 leu2Δ0 lys2Δ0 met15Δ0 trp1Δ63 ura3Δ0 med14ΔORF::KanMX pGPD1-OSTIR::HIS3 MED10-3xFlag-MNase::URA3* pSH1933 (ars cen *TRP1 pRGR1-med14Δ*TID) pSH1940 (ars cen *LEU2 pRGR1-med14 TID-3xV5-degron)* | This work |
| RDY1 | *BDF1*-degron, *bdf2Δ*, *RPB3*-Flag | *RPB3-3x Flag::NatMX, pGPD1-OSTIR::HIS3, bdf2Δ::HPH, BDF1-3xV5 IAA7::KanMX* | (Donczew and Hahn, 2021) |
| RDY33 | *BDF1*-MNase, *BDF2*-degron, *RPB3*-Flag | *RPB3-3x Flag::NatMX, pGPD1-OSTIR::HIS3, BDF2-3xV5 IAA7, BDF1-MNase::TRP1* | (Donczew and Hahn, 2021) |

**Table S4 References:**

Brachmann, C.B., Davies, A., Cost, G.J., Caputo, E., Li, J., Hieter, P., and Boeke, J.D. (1998). Designer deletion strains derived from Saccharomyces cerevisiae S288C: A useful set of strains and plasmids for PCR-mediated gene disruption and other applications. Yeast *14*, 115–132.

Cam, H.P., Noma, K., Ebina, H., Levin, H.L., and Grewal, S.I.S. (2008). Host genome surveillance for retrotransposons by transposon-derived proteins. Nature *451*, 431–436.

Donczew, R., and Hahn, S. (2021). BET family members Bdf1/2 modulate global transcription initiation and elongation in Saccharomyces cerevisiae. Elife *10*, e69619.

Donczew, R., Warfield, L., Pacheco, D., Erijman, A., and Hahn, S. (2020). Two roles for the yeast transcription coactivator SAGA and a set of genes redundantly regulated by TFIID and SAGA. Elife *9*, e50109.

Reeves, W.M., and Hahn, S. (2003). Activator-independent functions of the yeast mediator sin4 complex in preinitiation complex formation and transcription reinitiation. Molecular and Cellular Biology *23*, 349–358.
